# Minimal Mimics and Maps of Natural Light for Mammals

**DOI:** 10.64898/2026.01.18.698509

**Authors:** Philippe Morquette, Michael Tri H. Do

## Abstract

Light drives processes that include perception and the regulation of circadian rhythms, sleep, metabolism, and development. These processes are initiated by photopigment molecules, each preferentially absorbing particular wavelengths. Light of a given spectrum stimulates an animal’s set of photopigments in a specific profile. Natural skylight and its variations produce stimulation profiles that promote normal physiology. To mimic these profiles using artificial light, we consider the thermally stable, photoconvertible states of relevant photopigments: ground states of rhodopsin and cone photopigments, and three states of melanopsin. This gives a relatively high-dimensional representation of illumination. Nevertheless, we find that two wavelengths suffice to closely mimic the effects of natural light for mammals, including humans and mice. Adjusting the wavelength ratio allows mimicry of natural light’s variations, such as those from twilight to noon. Ratio adjustments also compensate for light’s filtering by elements like the eye’s optics and laboratory cages. Adding a third wavelength makes natural light mimicry nearly perfect. By contrast, common artificial lighting—designed for low-dimensional, human color space—stimulates photopigments in unnatural proportions. We conclude by providing species-specific maps of photopigment stimulation profiles under natural and artificial illumination, which make our observations intuitive while providing insight into the diverse visual ecologies of mammals.

**SIGNIFICANCE STATEMENT:** Humans sense light for vital processes like sight and physiological regulation. These processes are normal under natural skylight, where they evolved. However, much of modern life is spent in artificial light, which is unlike natural light in many of its biological effects. This disparity has been linked to disorders that range from cardiovascular disease to cancer. This manuscript introduces simple forms of artificial lighting that replicate the effects of natural skylight on photoreceptors of humans and other species. It also demonstrates how the biological effects of natural and artificial lights can be captured in simple maps, facilitating the choice and further design of illumination that is beneficial.

## INTRODUCTION

Humans and other animals use light to see and to regulate processes that include development, cardiovascular function, circadian rhythms, sleep, metabolism, cognition, and mood (Lucas et al., 2014; Do, 2019). Light is sensed by photopigments, G protein coupled receptors (opsins) bound to chromophore. Three types are known to trigger electrical signals: rhodopsin, cone photopigments, and melanopsin (Do, 2019; D’Souza et al., 2022). Each absorbs light of particular wavelengths between ultraviolet and near-infrared. Thus, various spectra of illumination produce specific profiles of photopigment stimulation. These profiles differ among species whose photopigments vary in their absorption spectra.

Mammalian photopigments evolved under ambient illumination from the sun. The spectrum and intensity of this natural skylight support normal biological processes. However, most light in modern society is artificial, which tends to be unnatural in its spectrum, too bright at night, and too dim during the day (Windred et al., 2024; Harmsen et al., 2025). Even natural light, when passed through windows and other materials, usually becomes unnatural. Abnormal lighting impairs the development of retinal structure and circadian period, increases mortality from cardiovascular events, dysregulates mood, and misaligns circadian rhythms to raise the risk of disorders such as cancer (Sahar and Sassone-Corsi, 2009; Lazzerini Ospri et al., 2017; Do, 2019; Windred et al., 2024). Conversely, light is therapeutic when properly designed and controlled (Voysey et al., 2021; Wescott et al., 2025). For these reasons, it is important to develop artificial light that stimulates photoreceptive mechanisms naturally.

Artificial light of this kind would produce the same photopigment stimulation profile as natural light. Most artificial lights do not. They may appear natural, being designed for human color perception and its underlying cone photopigments. However, rhodopsin is generally ignored; it has been thought to operate only in dim light, but its role is likely broader (Reinhard et al., 2025). Melanopsin’s importance is recognized in a new lighting standard (CIE, 2018a).

However, this standard concerns only one melanopsin state. Melanopsin has three stable states, which are important to consider for two main reasons (Matsuyama et al., 2012; Emanuel and Do, 2015; Emanuel and Do, 2023; Liu et al., 2023). First, their spectral separation is substantial, approaching that of the cone photopigments mediating the red-green axis of color vision. Second, they are photoconvertible—light switches the molecule between active and inactive states, fundamentally shaping its actions. Thus, a photostimulation profile of six photopigment states should be considered for dichromatic mammals (two cone photopigments, rhodopsin, and three melanopsin states) and seven for trichromatic mammals (adding a cone photopigment).

A natural light mimic should be able to compensate for elements in its path to the photopigments. These prereceptoral filters include window materials, lamp coverings, and the eye’s optics. Their effects can be strong. For example, aging of the human lens reduces transmission, especially of short wavelengths, potentially contributing to circadian and sleep dysregulation (Eto and Higuchi, 2023). Most laboratory cages attenuate ultraviolet wavelengths that mice and other species sense for perception and physiological regulation (van Oosterhout et al., 2012). Conversely, even if daylight were given directly to the retina, as in *ex vivo* physiological studies, it would act differently and potentially unnaturally due to the absence of filtering by the eye. The inability to deliver the same, naturalistic stimulus *ex vivo* and *in vivo* impedes efforts to connect biological mechanisms to behavior. Compensation is challenging because most light sources and prereceptoral filters have complex spectra.

We sought to define how natural light—defined here as skylight illumination at nautical twilight and above—stimulates rhodopsin, cone photopigments, and melanopsin’s three states for various species. We then asked if these photopigment stimulation profiles could be mimicked with direct artificial light that comprises few wavelengths and is thus readily corrected for diverse forms of prereceptoral filtering. Finally, we sought compact descriptions of photopigment stimulation profiles across variations in natural light, to facilitate studies of photoreception in ecological contexts as well as improvements in lighting design.

## MATERIALS AND METHODS

All experiments were conducted according to the protocols approved by the Institutional Animal Care and Use Committee at Boston Children’s Hospital.

### Materials Availability

This study did not generate new unique reagents.

### Data and Code Availability

All study data are included in the article and/or Supplementary Material. All data are available upon request from Philippe Morquette.

### Estimating photopigment stimulation profiles in natural and artificial light

All spectra were expressed in units of photon flux density across wavelength (photons µm^-2^ s^-1^ nm^-1^). Each light spectrum was measured for this study, published previously, or is a standard. The light spectrum was multiplied by the normalized transmission spectrum of each prereceptoral filter considered. The product was multiplied with the spectral sensitivity of a photopigment, then integrated to give the total photons available to stimulate the photopigment (Rushton, 1972). For brevity, this integral is referred to as the photopigment’s “stimulation.” The spectral sensitivity of a photopigment was given by its λ_max_ and the mathematical template (nomogram) of Govardovskii and colleagues (Govardovskii et al., 2000), which includes the main α band and the secondary β band. The γ band was not considered because it is a non-specific absorption peak that is common to most proteins, not specified in existing nomograms, and excites the photopigment modestly; furthermore, photons at these wavelengths are sparse in natural light spectra (Fein and Szuts, 1982; Hernandez-Andres et al., 2001; Johnsen et al., 2006; Spitschan et al., 2016; ASTM, 2023).

Repeating this procedure for each relevant photopigment state produced a series of values. For example, in mice, six photopigment states were considered: the ground states of rhodopsin (Opn2) and of the short- and medium-wavelength-sensitive cone photopigments (Opn1SW and Opn1MW), and the three known states of melanopsin (Opn4-R, -M, and -E). These photopigment stimulations were combined into a six-element vector, which was converted into a unit vector to compare spectra of different intensities. For brevity, this vector is referred to as the “photopigment stimulation profile” or simply “profile” of a natural or artificial light. Each stimulation profile is specific to one species’ complement of photopigments under one light spectrum and one condition of prereceptoral filtering.

### Photopigment self-screening and co-screening

Photopigment self-screening was accounted for in humans and mice, for which data are available. Each photopigment’s absorption spectrum was adjusted using this equation (CIE, 2006):

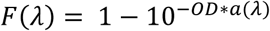

where a(λ) is the photopigment’s Govardovskii nomogram (normalized) and OD is its axial optical density in the phototransducing outer segment (OS). The corrected spectrum was normalized by its peak. Self-screening was omitted for melanopsin because of its low expression density in ipRGCs (Do et al., 2009).

For the human, OD values were 0.35 for Opn2 (Alpern and Pugh, 1974), 0.30 for Opn1SW, and 0.38 for Opn1MW and Opn1LW (CIE, 2006).

For the mouse, the OD of Opn2 was 0.456 (Naarendorp et al., 2010). Accounting for Opn1SW and Opn1MW was more complex, as most cones co-express them in reciprocal gradients across the retina (Wang et al., 2011). Thus, in single cones, these photopigments screen themselves and one another to different degrees. A simplification was used in which individual cones were considered to express either Opn1SW or Opn1MW alone, each with an OD of 0.195, computed as 0.015 OD µm⁻¹ × 13.0 µm (the OS length)(Lyubarsky et al., 1999).

To assess the fairness of this simplification, mutual screening was assessed. Opn1SW and Opn1MW were considered to be intermixed in the OS. The total absorbance of the OS was

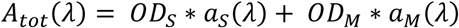

where *OD_S_* and *OD_M_* are the optical densities of Opn1SW and Opn1MW, respectively. *aS(λ)* and *aM(λ)* are Govardovskii templates for their absorption spectra.

From the Beer–Lambert law, the total fraction of incident photons absorbed is

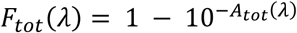

Each photopigment absorbed photons in proportion to its relative absorbance at each wavelength. The mutually-screened photopigment absorption spectra are

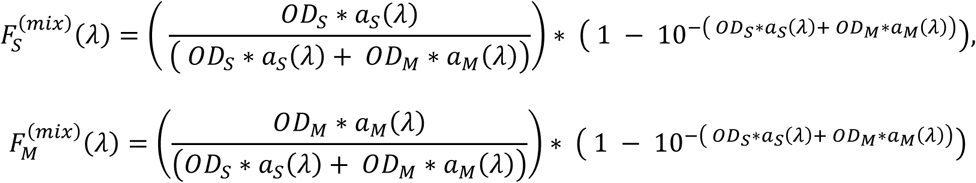

The most extreme case is for Opn1MW within a cone containing 99% Opn1SW (Wang et al., 2011). Under the 2-wavelength mimic of the G173 daylight standard (ASTM, 2023), stimulation of this mutually-screened Opn1MW deviates from that of Opn1MW (self-screening only) by - 3.5%. For Opn1SW, its smallest proportion is within a cone containing 65% Opn1MW (Wang et al., 2011). Under the 2-wavelength G173 mimic, stimulation of this mutually-screened Opn1SW deviates from that of Opn1SW (self-screening only) by −10.9%. A case was then considered where Opn1SW and Opn1MW are present in equal proportions. Under the 2-wavelength G173 mimic, stimulation of these mutually-screened photopigments deviates from those with self-screening by 8.5 and −1.8%, respectively, resulting in an overall relative root mean squared error (RRMSE) across all photopigments of 3.4%. Mutual screening thus appears to modestly affect the accuracy of natural light mimicry.

### Transmission spectrum of the mouse eye

Male and female adult (>P56) C57BL/6J mice were deeply anesthetized with intraperitoneal Avertin and their pupils dilated with tropicamide (0.5-1%). Their eyes were removed into Ames’ medium (US Biological or Millipore Sigma) that was supplemented with 1.8 g/L of NaHCO_3_ and equilibrated with 95% O_2_/5% CO_2_ to achieve a pH of 7.4. The posterior pole of the eye was cut away near the ora serrata to open a light path through the vitreous, lens, aqueous, and cornea. The eye was mounted in a quartz cuvette (200-2500 nm transmission, 3.5-ml volume) filled with HEPES-buffered Ames’ medium at 35 °C (in mM, 140 NaCl, 3.1 KCl, 0.5 KH_2_PO_4_, 1.2 CaCl_2_, 1.2 MgSO_4_, 6 glucose, and 10 HEPES; pH adjusted to 7.4 with NaOH)(Do et al., 2009; Emanuel et al., 2017; Milner and Do, 2017). The transmission spectrum (300–700 nm) was obtained using a Shimadzu 1800 UV-Vis spectrophotometer. The background was subtracted using the same cuvette with no eye. For each eye, 5-10 spectra were averaged, ensuring no systematic change in the spectra over time. These average transmission spectra were consistent across eyes when acquired immediately after dissection (<5 min; transmission decreased, primarily over shorter wavelengths, over time). The ocular transmission spectrum showed a longpass quality, declining by 18.1 ± 0.6% from 700 to 400 nm (11 eyes, mean ± SD). Nevertheless, transmission of shorter wavelengths is greater than previously reported (Henriksson et al., 2010).

### Determining optimal wavelengths and intensities for natural light mimicry

To design mimics of natural light’s effects on the photopigments of a given species, a matrix was constructed with each row corresponding to a photopigment and each column to a wavelength. Thus, for the mouse 2-wavelength mimic, the matrix was 6×2. Wavelengths were selected randomly within a defined range (e.g., 380-700 nm for the mouse). To determine the intensities of each wavelength that would provide the closest match of photopigment stimulations to natural light, an error function was defined:

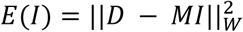

Where E is the error, D is the normalized (unit norm) photopigment stimulation profile under the daylight spectrum in question (e.g., G173), and the mimic profile is the vector 𝐷^ = 𝑀𝐼 generated by a given wavelength combination and its intensities. *M* is the matrix that captures the stimulation profiles at intensity 1 for a given set of wavelengths and *I* is a vector containing the intensities of each wavelength. The weights (W) are the inverse square of each photopigment’s target stimulation profile value, ensuring that error contributions are relative and no photopigment dominates the fit. The optimal intensity vector (*I_opt_*) that minimizes E is derived analytically as

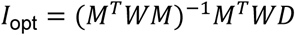

Using 𝐼_opt_, the predicted profile 𝐷^ = 𝑀𝐼_opt_ is computed and then normalized (to unit norm) before being compared to 𝐷 using

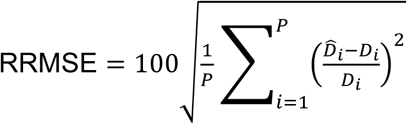

where 𝑃 is the number of photopigments. Each component was divided by 𝐷_𝑖_to express the error for each photopigment in relative (fractional) units rather than absolute units. This makes RRMSE a component-wise relative RMSE that matches the weighting 𝑊 = diag(1/𝐷_i_^2^) used in 𝐸(𝐼).

This process was repeated 1000-10,000 times, discarding any *I* containing negative values, and the ten wavelength combinations with the lowest RRMSE were selected for further optimization using gradient descent (using the L-BFGS-B algorithm)(Zhu et al., 1997), which further refined the wavelengths (with intensities determined analytically as *I_opt_* for each candidate wavelength set). Each intensity is the integral of the light’s spectrum (in units of photons). Mimic accuracy depends on the ratio of intensities. The overall intensity (keeping the ratio constant) may be set to the desired level.

### Determining practical ranges of wavelength and intensity for natural light mimicry

Because real light sources have spectra of discrete widths, it is useful to identify ranges of wavelengths that support adequate natural light mimicry. For a 2-wavelength mimic, a matrix was constructed whose dimensions are the first stimulus wavelength (λ1), the second stimulus wavelength (λ2), the ratio of their intensities (λ ratio = intensity λ1 / intensity λ2), and the complement of photopigments (i.e., six additional dimensions in mouse). The wavelengths were from 300–700 nm at 0.2-nm resolution, and the ratio ranged from 0.1 to 10 in increments of 0.01. Each value in the matrix is a photopigment’s stimulation by two wavelengths at a given ratio. Each profile (e.g., the vector containing stimulation of each photopigment, for a given choice of λ1, λ2, and λ ratio) was compared to that of the reference spectrum (e.g., D65 for humans) to generate an RRMSE. This operation was performed for all vectors, resulting in a 3-dimensional matrix (λ1, λ2, and λ ratio) of RRMSEs. This matrix was used to generate a color map, in which parameters producing an RRMSE <10% were highlighted. The 3-wavelength mimic had two ratios, λ ratio_1_ (intensity λ1 / intensity λ3) and λ ratio_2_ (intensity λ2 / intensity λ3).

### Construction of natural light mimics for humans

A Ganzfeld diffusing sphere was constructed from a hollow acrylic globe of 30.5-cm diameter (Crown Plastics, 20012-SM-5N). A smaller version was built using 2 aluminum hemispheres of 12.7-cm diameter (Wagner, B4162). The larger globe had a 13-cm opening that served as the viewing port, covering ∼130-140° of visual angle (>80% of a human eye’s visual field)(Strasburger et al., 2011). The smaller Ganzfeld had a viewing port of 3.4 cm. For both spheres, a second, smaller port was added to accommodate a liquid light guide (5-mm core, Thorlabs, LLG05-4H). This port was positioned such that direct illumination from the light guide was outside the field of view; all viewed light was internally reflected. For diffuse internal reflection that is even across wavelengths, the globes were internally coated with layers of BaSO_4_ until uniform and opaque (Edmund Optics, 83-890).

*2- wavelength D65 mimic.* Light from LEDs (Solis-470C and Solis-565D) was conditioned with bandpass filters (FBH460-10 and FL457.9-10 in series for the short-wavelength beam, and FBH550-10 for the medium-wavelength beam), and combined using a beamsplitter (DMSP490R; all parts from Thorlabs).
*3- wavelength D65 mimic.* Light from LEDs (Solis-470C, Solis-525C and Solis-565D) was conditioned with bandpass filters (FBH455-10, FBH520-10 and FBH590-10, respectively) and combined with beamsplitters (DMSP490R and DMSP550R; all parts from Thorlabs).

For both mimics, the combined beam was focused on the end of the liquid light guide using an aspheric lens. Power was controlled by current level (DC20, Thorlabs). The output of each Ganzfeld sphere was measured using a radiometer (PM100D and S130VC, Thorlabs) and spectrometer (Flame-T-UV-VIS, Ocean Optics; coupled to a 400-µm diameter fiber). The sensors were placed directly in front of the viewing apertures for measurement. The power of each LED was adjusted until the desired spectrum was obtained.

### Human perception of natural light mimics

To mimic natural light for both foveal and peripheral vision, we used the CIE 1964 color matching functions (CMFs; 10° standard observer covering a large field around the fovea)(CIE, 2019b). Following standard procedure, the power spectral density of the mimic was multiplied by the corresponding x̂, ŷ, and ẑ values from the CMFs, and the products integrated to obtain the raw X, Y, and Z tristimulus values; the normalization constant was set such that Y=100.

From these normalized values, the chromaticity coordinates (x, y) were computed by dividing each of X, Y, and Z by their sum (i.e., X+Y+Z)(CIE, 2018b). The mimics and D65 coordinates were then projected onto a CIE 1964 chromaticity diagram. For screen visualization, a regular grid of (x, y) coordinates within the diagram was linearly transformed into sRGB color space, and gamma correction (IEC, 2003) was applied to align the rendered colors with human perception.

To quantify the color differences between the D65 illuminant and its 2- and 3-wavelength mimics, their XYZ tristimulus values were converted into the CIELAB color space, and the CIEDE2000 formula was used to estimate their color difference (ΔE_00_) under the Ganzfeld viewing conditions (Luo et al., 2001; CIE, 2018b).

### Maps of photopigment stimulation profiles in natural light

Photopigment stimulation profiles (unit vectors) were computed for 2600 natural light spectra measured at solar angles of −8-76° in Granada, Spain (only 56 spectra below 0°), and for 967 natural light spectra measured at solar angles of −13-24° (744 spectra below 0°) in rural Pennsylvania, USA (Hernandez-Andres et al., 2001; Spitschan et al., 2016). These were the most complete, readily available sets of natural light that met inclusion criteria (range of at least 300–700 nm, resolution of ≤5 nm, and more than one measurement at a given solar elevation in a given terrestrial location).

Principal component analysis (PCA) was performed on a matrix containing the 3,567 profiles (using the scikit-learn package in Python), where each row corresponded to a profile obtained for a spectrum and each column to one photopigment’s stimulation (see above) in the set of photopigments present in a species. The first two principal components were used for visualization.

### Cluster analysis

DBSCAN, a density-based clustering algorithm, was used to assess the closeness of natural and artificial photopigment stimulation profiles (Ester et al., 1996; Pedregosa, 2011). It was applied to all dimensions of these profiles (e.g. 6 for the dichromatic mouse). DBSCAN is advantageous because it does not require specifying the number of clusters and can detect clusters of arbitrary shape. Furthermore, its ability to classify low-density points as noise makes it robust to outliers, which is ideal when comparing diverse natural and artificial light sources.

HDBSCAN was also considered but found to produce clusters that were not robust to small changes in clustering parameters or to subsampling of the data set.

The parameters of DBSCAN are MinPts, the minimum number of neighbors that a point must have to be the core of a cluster, and Eps, the maximum distance over which a point can be considered a neighbor. These parameters were initialized using heuristics that are based, for transparency and reproducibility, solely on the properties of the data (Sander et al., 1998; Schubert et al., 2017). MinPts was set to 2*dimensions. Eps was first set based on the distance to the k^th^ nearest neighbor (kNN), where k = (2*dimension-1); for 6-dimensional mouse data, this parameter is 11. For each data point, the kNN distance was then calculated. Data points were ordered by their kNN distance and plotted as a k-distance graph. The value at the point of maximum curvature, where the k-distance begins to increase steeply, was identified as the updated Eps. Points with kNN distances smaller than the updated Eps were assigned to a cluster, and others considered noise (Sander et al., 1998; Schubert et al., 2017). Most of the 3,567 natural stimulation profiles were within a single cluster, with <1.3% excluded.

Bootstrapping was used to ask if artificial profiles clustered with the natural profiles. In each iteration, 80% of the 3K profiles were randomly sampled. An artificial profile was then added to the resampled data and DBSCAN applied (maintaining the Eps and MinPts values determined for the full data set). This process was repeated for 1,000 iterations, with a new resampling for each. Resampling was performed without replacement to mitigate bias from varied local densities of natural profiles. If an artificial profile clustered with the natural profiles in ≥95% of iterations, it was classified as being similar.

## RESULTS

We will start by explaining our process for estimating spectra for natural light, ocular transmission, and photopigment absorption. We will use these spectra to design natural skylight mimics using two or three wavelengths delivered in the correct ratio for mice, rats, cats, rabbits, dogs, and humans. We then compare the accuracy of wavelength mimics in theory and practice for humans. Throughout, we provide simple maps of photopigment stimulation profiles under natural and artificial direct illumination. We will focus our analyses on photopigments that are known to trigger electrical responses in the mammalian retina, those of rods (rhodopsin), cones (cone photopigments), and intrinsically photosensitive retinal ganglion cells (ipRGCs; melanopsin).

### Choosing photopigment absorption spectra

We consider photopigment states that are stable and thus likely to absorb a photon and produce a biological effect (i.e., we ignore short-lived intermediate states). For rhodopsin and cone photopigments, which are monostable, this is only the ground state; following activation, the opsin loses its chromophore and becomes insensitive to light (Luo et al., 2008). Mammalian melanopsin is multistable (Matsuyama et al., 2012; Emanuel and Do, 2015; Emanuel and Do, 2023; Liu et al., 2023). Its three known conformations are melanopsin (the R state, non-signaling), extramelanopsin (E, non-signaling), and metamelanopsin (M, signaling). Photon absorption by R or E can produce M, while absorption by M can produce R or E. Melanopsin’s actions depend on the dynamic photoequilibration of these states.

Several absorption spectra have been reported for each photopigment state, with varying wavelengths of peak sensitivity (λ_max_). We prioritize *in vitro* measures because they are direct and choose λ_max_ values that are most consistent across spectrophotometry (purified photopigment), microspectrophotometry (photopigment in cells), and electrophysiology (the electrical activity of photopigment in cells)(**Table S1**). We use the mathematical function of Govardovskii and colleagues to reconstruct each photopigment state’s absorption spectrum, with only the λ_max_ as a free parameter (Govardovskii et al., 2000). These reconstructed spectra are highly accurate across log units of variation in absorbance, though complexities exist (Luo et al., 2011; Matsuyama et al., 2012; Emanuel and Do, 2023). We also corrected absorption spectra for self-screening of photopigments in mouse and human photoreceptors (Alpern et al., 1987; CIE, 2006).

### Mimicking natural light for mouse photopigments

We begin with the mouse because of its widespread use as a model system and detailed knowledge of its photoreceptive mechanisms. We use photopigment λ_max_ values of 502 (rhodopsin, Opn2)(Sakurai et al., 2007), 358 (short-wavelength-sensitive cone photopigment, Opn1SW)(Tsutsui et al., 2007) and 508 nm (medium-wavelength-sensitive cone photopigment, Opn1MW)(Sakurai et al., 2007; Tsutsui et al., 2007). For the R, E, and M states of melanopsin (Opn4-R, Opn4-E, and Opn4-M), these values are 471, 453, and 476 nm, respectively (Matsuyama et al., 2012; Emanuel and Do, 2015; Emanuel and Do, 2023). We take the R and E state λ_max_ values from electrophysiological measurements of mouse ipRGCs, as they concern melanopsin in its native environment (Emanuel and Do, 2015). These values are slightly red-shifted from those of purified mouse melanopsin (467 and 446 nm, respectively)(Matsuyama et al., 2012). The M state’s λ_max_ is only known for purified melanopsin, so we use it as is (i.e., without imposing a red shift that may be inferred from electrophysiological assessments of the other states).

The light that reaches the retinal photopigments is shaped by the transmission spectrum of the eye, primarily the cornea and lens. A spectrum has been published for the mouse eye but only for a limited wavelength range (Henriksson et al., 2010). Thus, we dissected the mouse eye, removed a portion of the posterior pole, and mounted the remainder in a custom cuvette for spectrophotometry (**Materials and Methods**). We worked swiftly and used oxygenated media to maintain tissue health. Compared to the published spectrum, ours has higher transmission of short wavelengths (**Fig. S1**).

Multiplying light’s spectrum (photons µm^-2^ s^-1^ nm^-1^), the ocular transmission spectrum (normalized to its peak), and a photopigment state’s absorption spectrum (normalized to its peak) yields the light that is available to that state (**Fig. 1*A***). Downstream factors—for instance, the light that the photopigment actually absorbs, the probability that the photopigment activates, the probability that activated photopigment triggers its G protein cascade, properties of the cascade, and biological effects that may vary across retinal subregions—follow from photon availability (**Materials and Methods**)(Polyak, 1941; Rodieck, 1998; Johnsen, 2012; Mouland et al., 2019; Stockman and Rider, 2023). Integrating the product gives the total photon flux density that is available for the photopigment state in question (**Fig. 1*B***). For brevity, we refer to this integral as a photopigment’s stimulation.

**Figure 1.**
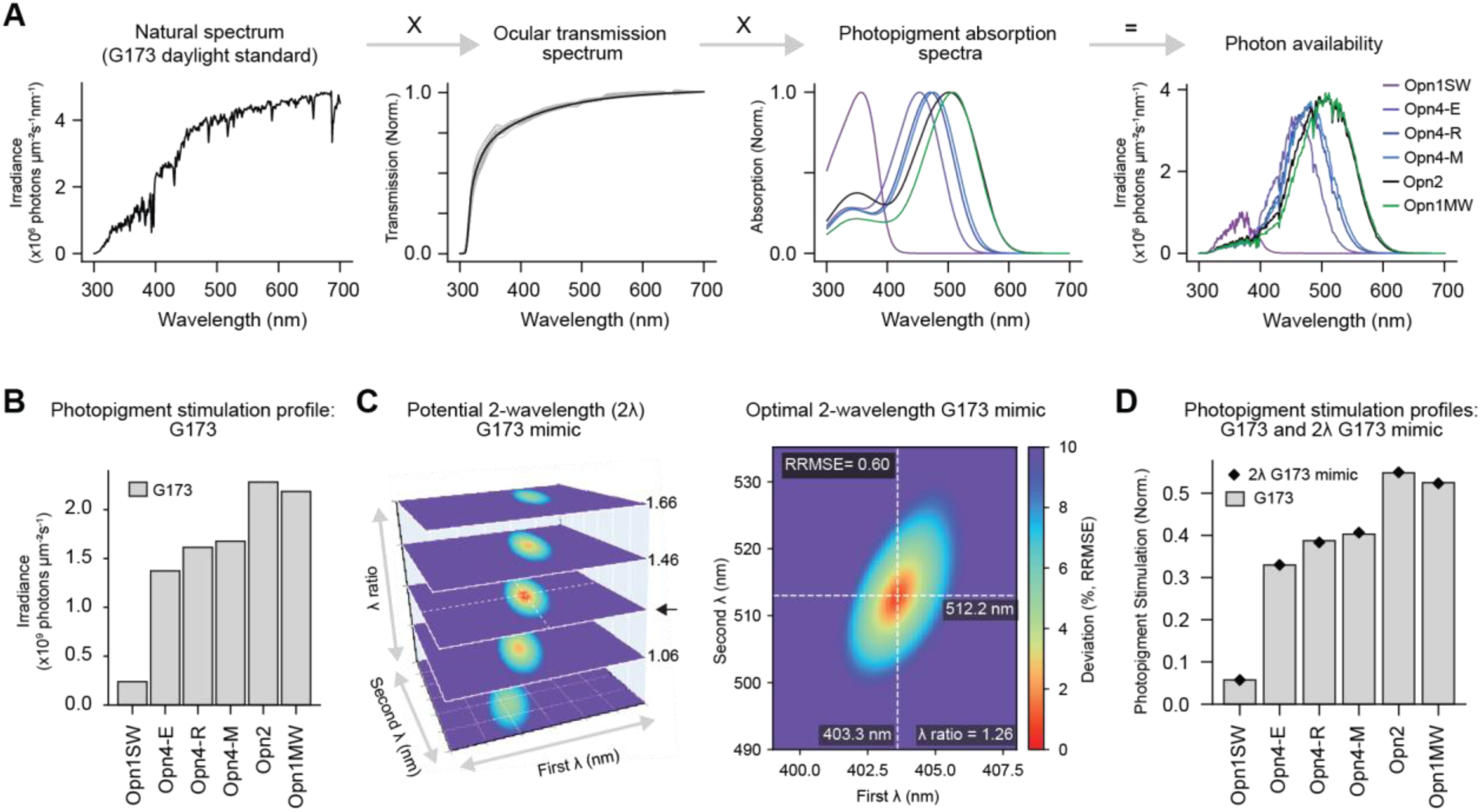
**2-wavelength mimicry of natural light for a dichromatic mammal (mouse). *A***, Multiplying the spectrum of a daylight standard (ASTM G173) by the normalized transmission spectrum of the mouse eye (see **Fig. S1**) and the normalized absorption spectrum of each mouse photopigment yields the photon availability for each photopigment (or, in the case of melanopsin, photopigment state). ***B***, Integrating each photon availability spectrum gives the G173 photopigment stimulation profile (“profile”). Photopigments are ordered by peak wavelength sensitivity, from shortest to longest. The three stable states of melanopsin (Opn4) are given individually. ***C***, Varying two wavelengths (first and second λ) and their ratio (vertical axis) reproduces the G173 profile with varying accuracy (the deviation is given as a relative root mean squared error, RRMSE, across photopigments). The G173 profile is mimicked optimally by wavelengths of 403.3 nm and 512.2 nm, given in a ratio of 1.26:1. The color code for deviation is shown on the right. ***D***, The profiles of G173 (gray bars) and its 2-wavelength mimic (black diamonds). Profiles are expressed in terms of their unit vectors to focus on spectrum rather than intensity.

We first consider a standard daylight spectrum, ASTM International’s G173-tilted (ASTM, 2023), which captures both direct and diffuse illumination at mid-day under a cloudless sky. We examined the range from 300 nm (photon flux is scarce at shorter wavelengths, especially after attenuation by the cornea and lens) to 700 nm (at which point the mouse photopigment sensitive to the longest wavelength only absorbs at ∼10^-5^ of its peak) at a resolution of <1 nm.

The G173 spectrum, with consideration of the eye’s optics and each photopigment’s absorption, stimulates Opn1SW, Opn4-E, Opn4-R, Opn4-M, Opn2, and Opn1MW photopigments at 2.38×10^8^, 1.37×10^9^, 1.61×10^9^, 1.68×10^9^, 2.28×10^9^, and 2.19×10^9^ photons µm^-2^ s^-1^, respectively (**Fig. 1*B***; we order photopigments from shortest to longest λ_max_). This photopigment stimulation profile reveals a nearly 10-fold difference in skylight’s stimulation across the mouse photopigments.

Inspection of the mouse’s photopigment absorption spectra (**Fig. 1*A***) suggests that only two wavelengths may suffice to mimic the photopigment stimulation profile under skylight. That is because one wavelength can be chosen to intersect most absorption spectra at various levels, and a second wavelength can provide fine tuning. To test this idea, we searched for a pair of wavelengths that produce an artificial profile with the lowest deviation from the G173 profile (**Materials and Methods**). We focused on the spectrum of light rather than its intensity by normalizing the profiles of G173 and potential 2-wavelength mimics (i.e., comparing their unit vectors). We found that optimal wavelengths were 403.3 and 512.2 nm, given at an intensity ratio of 1.26:1 (in units of photons; **Fig. 1*C*-*D***). This 2-wavelength G173 mimic has a stimulation profile that deviates <1.1% from that of G173 for each photopigment; averaged over all photopigments, its deviation is 0.60% (relative root mean squared error, RRMSE). The overall intensity of the two wavelengths can be adjusted to match the photon flux density of G173 or any desired illumination level, from the threshold of perception to maximum daylight.

### Mimicking natural light’s variations for mouse photopigments

To ask if two wavelengths can mimic natural light’s variations, we compiled a large number of published spectra. We chose spectra that have a range of at least 300 to 700 nm (see above). We also selected for a minimum resolution of 5 nm. A ≤10-nm band is often considered monochromatic in vision studies. Extending the wavelength range or resolution would greatly reduce the available spectra. We arrived at a set of 3,567 spectra measured in Pennsylvania, USA (Spitschan et al., 2016) and Granada, Spain (Hernandez-Andres et al., 2001). Together, they cover solar elevations from nautical twilight to bright daylight (**Fig. 2*A***). These spectra are in units of photon flux density across wavelength (photons µm^-2^ s^-1^ nm^-1^)(Johnsen, 2012). We multiplied each spectrum by those of the mouse eye and its photopigments to yield a large set of photopigment stimulation profiles, which we refer to as the 3K profiles. Again, to focus on the spectrum rather than intensity of light, we converted each profile to its unit vector (**Fig. 2*B***).

**Figure 2.**
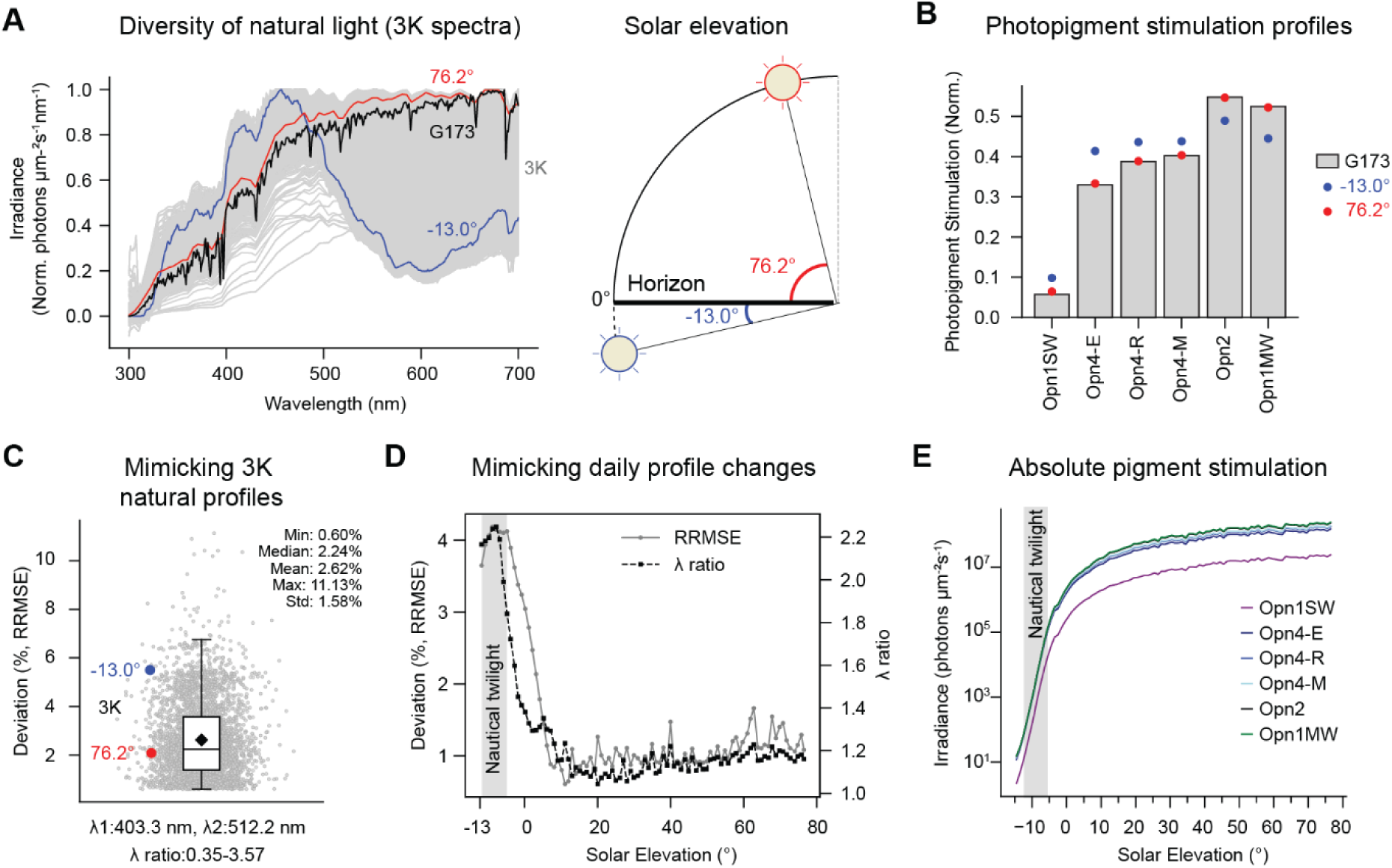
**2-wavelength mimicry of natural light’s variations for a dichromatic mammal (mouse). *A***, Spectra of the G173 daylight standard (black) and 3,567 (3K) samples of natural light (gray traces)(Hernandez-Andres et al., 2001; Spitschan et al., 2016). Highlighted are spectra taken at the lowest (−13°, blue) and highest (76.2°, red) solar elevations in the 3K set. ***B***, Photopigment stimulation profiles generated from the blue and red spectra in ‘***A***’ compared with that of G173. ***C***, The two wavelengths for optimal G173 mimicry (403.3 and 512.2 nm), with their ratio adjusted (0.35-3.57), mimic most 3K profiles (gray points) with <10% deviation. Points are spread along the x axis for visualization. Shown are the interquartile range (IQR, box), median (line), mean (diamond), and last point within 1.5×IQR. The −13° (blue) and 76.2° (red) solar elevations are highlighted. ***D***, Two wavelengths, optimized for an average trajectory of daily spectral variations (402.6 and 510.8 nm) with ratio adjustment (black), mimic that trajectory with little deviation (RRMSE, gray). Deviation is negligible at lower elevations where only rhodopsin requires consideration (gray box); the stimulation of one photopigment can be set precisely just using intensity. ***E***, Photopigment stimulation as a function of solar elevation, considering both the spectrum and intensity of illumination. The gray box indicates nautical twilight, below which only rhodopsin requires consideration.

We asked if the wavelengths we found for mimicking G173 (403.3 and 512.2 nm) could be used to mimic the 3K profiles. Adjusting the ratio of these wavelengths from 0.35 to 3.57 allowed mimicry of 90.1% of the 3K profiles with <5% deviation (**Fig. 2*C***). We also optimized the two wavelengths for all 3K profiles, obtaining values of 402.5 and 508.9 nm. Changing their intensity ratio suffices to mimic 98.0% of the 3K profiles with <5% deviation (**Fig. S3*A***). The overall intensity of these wavelengths can be adjusted to match any level of environmental illumination. Therefore, two wavelengths in the correct ratio largely suffice to mimic the spectrum of natural skylight as it varies across parameters that include geographical location and meteorological conditions.

Daily variations in the spectrum of natural skylight are salient cues for processes that include synchronization of the circadian clock with the day-night cycle, and are sensed in part by mechanisms that compare wavelengths (e.g., via color opponency)(Solessio and Engbretson, 1993; Dacey et al., 2005; Walmsley et al., 2015; Spitschan et al., 2017a; Mouland et al., 2019; Patterson et al., 2022; Liu et al., 2023). These spectral changes are more pronounced from astronomical to civil twilight, when the sun approaches the horizon (Spitschan et al., 2016).

Therefore, we assessed the effectiveness of a 2-wavelength mimic that is specifically designed to capture these variations. We pooled 3K profiles in 1° bins of solar elevation, from −13 to 76° (just below nautical twilight to the highest elevation in the 3K spectra). This gave the average trajectory of daily spectral variations. We found two optimal wavelengths, 402.6 and 510.8 nm which, given in ratios between 1.00 to 1.40, have <2% deviation from profiles of spectra measured at solar elevations above 0° (**Fig. 2*D***). Accuracy declined below 0°, to a maximum of 4.1% deviation. In practice, this decrease in accuracy has little consequence because it is most prominent when only rods are sensitive enough to drive photoreception (i.e. photon flux density lower than ∼200 photons μm^-2^ s^-1^ between 300 and 700 nm, corresponding to light intensities below nautical twilight, **Fig. 2*E***). In this regime, one only needs to consider how the spectrum of light stimulates rhodopsin, and the target stimulation level can be achieved simply by adjusting the intensity. Hence, two wavelengths given in varying proportions suffice to mimic natural skylight’s variations from twilight to full daylight for mouse photopigments.

Given the effectiveness of two wavelengths, we asked if three wavelengths provided a substantial improvement (**Fig. S5*D***). With three wavelengths, mimicry of the G173 profile appears virtually perfect, with RRMSE values of <1×10^-6^%. Three wavelengths also increased accuracy for mimicking any of the 3K natural light profiles (**Fig. S4*A***). Thus, we consider three wavelengths to be ideal for mimicking natural skylight, and two wavelengths to be both accurate and practical.

### Photopigment stimulation profiles of artificial lights

We investigated how artificial lights stimulate mouse photopigments. Given the vast variety of artificial lights, we began by examining 42 standard spectra that represent typical sources (CIE, 2018b). All produced photopigment stimulation profiles that deviated substantially from that of G173 (>25% RRMSE; **Table S2**). However, many of these standards lack shorter wavelengths; they were designed for human visual perception, which is negligible for those wavelengths due to strong prereceptoral filtering and the relatively red-shifted absorption spectra of cone photopigments (see below). Shorter wavelengths are relevant for mouse photoreception (**Fig. 1*A***)(van Oosterhout et al., 2012). Therefore, we measured a selection of xenon, LED, halogen, incandescent, fluorescent, and monochromatic (10-nm band) sources and calculated their photopigment stimulation profiles (**Fig. 3*A-B***). Xenon was closest to G173. Its maximum deviation, for Opn1SW, was 36%. All other artificial profiles had deviations of >71% for at least one photopigment (**Fig. 3*B-C***). Finally, we considered monochromatic lights in theoretical, 0.01-nm bands from 380-700 nm. Each of their profiles deviated from that of G173 by >97% for one or more photopigments. Substantial deviations are also evident when considering the average of all photopigment stimulations in each profile (RRMSE; **Fig. 3*D***). Thus, all artificial lights examined differed substantially from a natural skylight standard.

**Figure 3.**
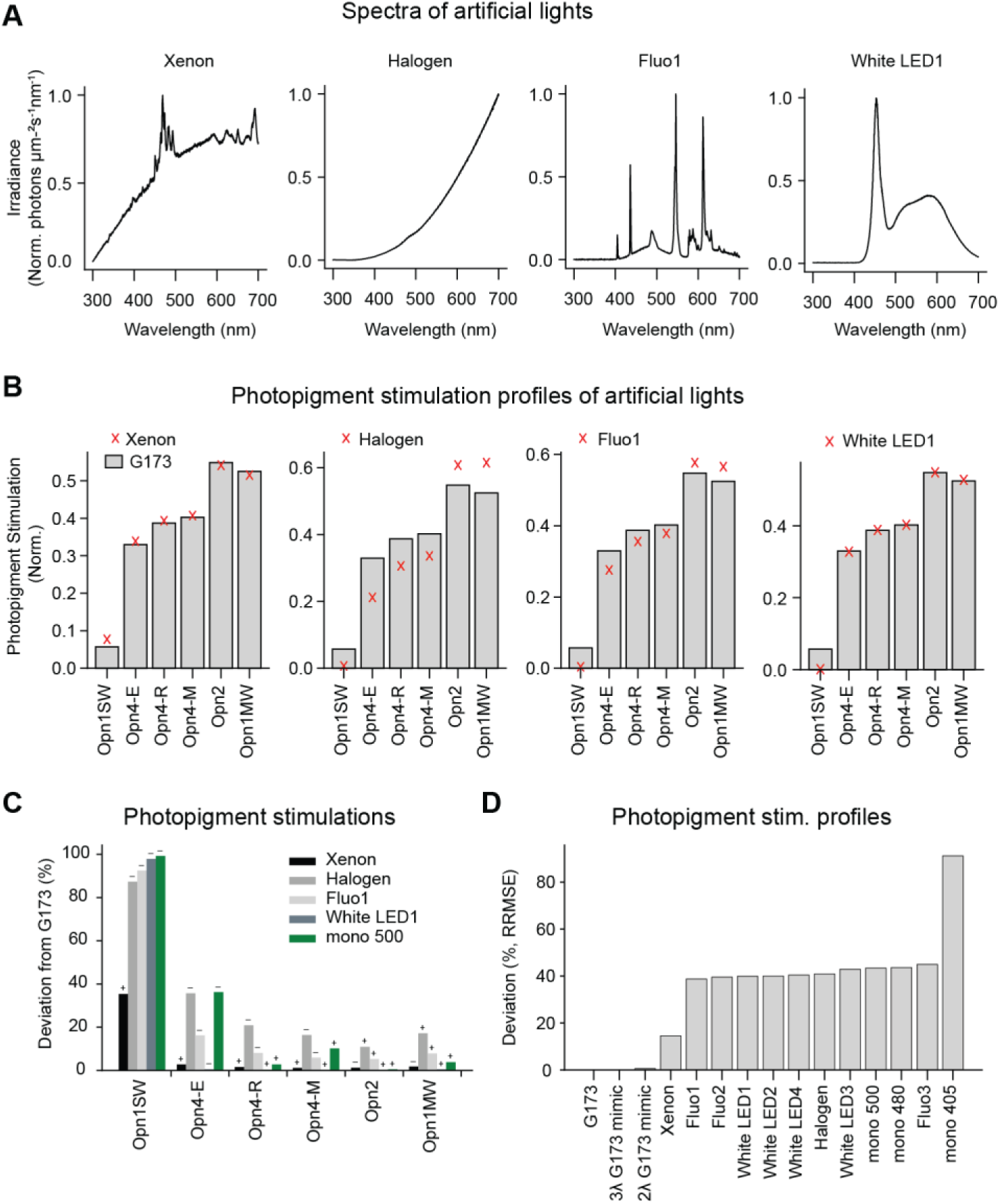
**Photopigment stimulation profiles of artificial lights. *A***, Example spectra measured from artificial lights. ***B***, Photopigment stimulation profiles from the artificial lights in ‘***A***’ (red x symbols) and the G173 standard daylight (gray bars). ***C***, Compared to G173, measured artificial lights produce higher (+) or lower (-) stimulation of individual photopigments. ***D***, Comparison of the G173 profile with those of measured artificial lights (deviation is expressed as the relative root mean square error, RRMSE, across photopigments). The 2- and 3-wavelength G173 mimics outperform all artificial lights, including broadband and monochromatic (‘mono’) sources.

### A map of mouse photopigment stimulation profiles

We found that the 3K natural light profiles were almost entirely captured in two dimensions, with 99.9% of their variance explained by the first two principal components (PC). Thus, the projection of the 3K profiles onto these PCs provides a simple and complete map of how natural light’s variations stimulate the mouse photopigments. Indeed, changes in solar elevation produce profiles that follow an orderly trajectory through the map (**Fig. 4*A***). Profiles of other available natural light spectra fell into the cloud formed by the 3K profiles (Johnsen et al., 2006)(not shown). The profiles of G173, and all aforementioned mimics were also within the cloud (**Fig. 4*B***).

**Figure 4.**
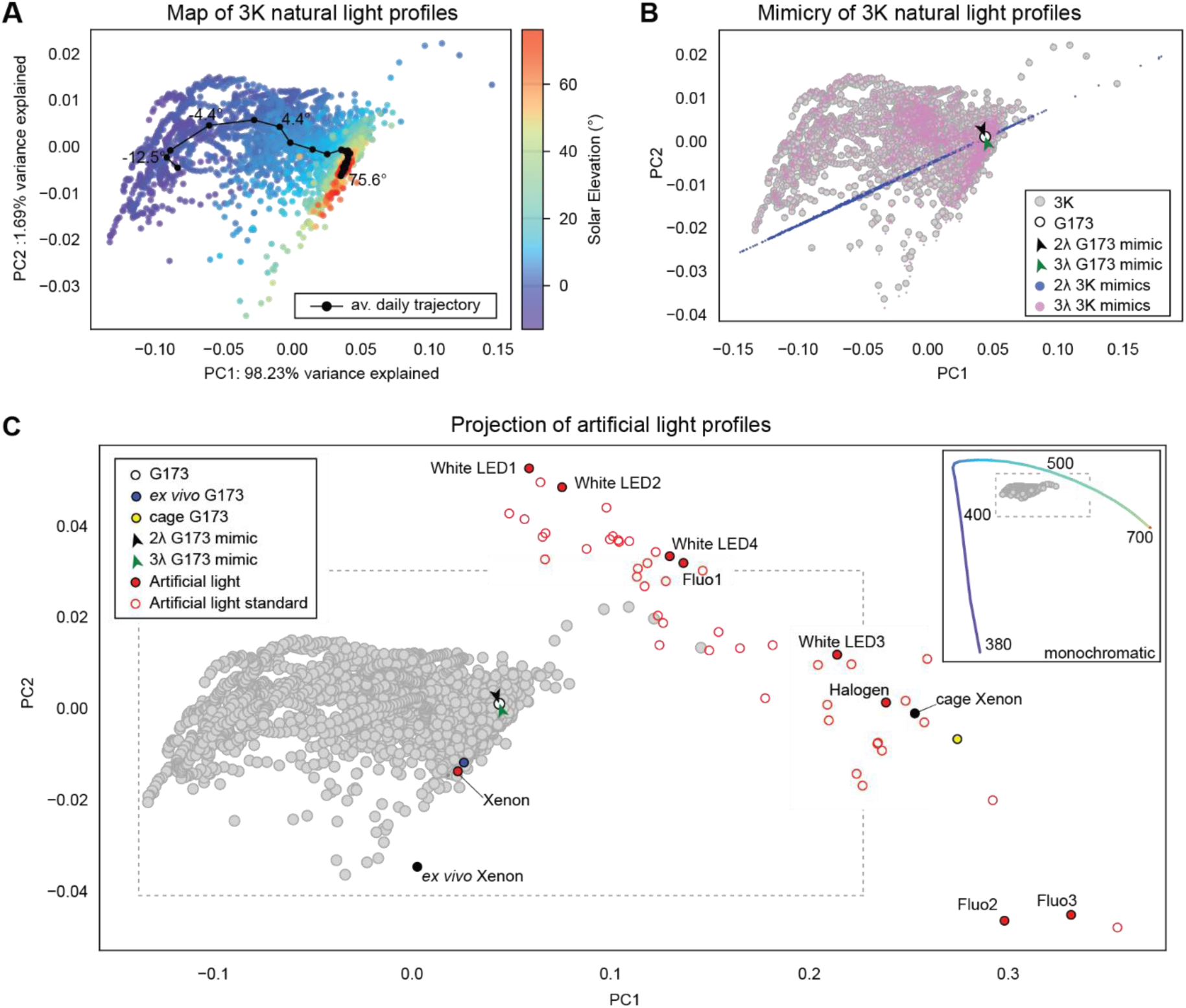
**Mapping photopigment stimulation profiles under natural and artificial lights for a dichromatic mammal (mouse). *A***, The 3K set of natural spectra produces mouse photopigment stimulation profiles (dots) that are well-described by their first two principal components (>99.9% of the variance explained). Profiles are color-coded by solar elevation, showing their systematic distribution within this 3K map. This distribution is also evident in the trajectory of profiles during an average day (black dots and line). ***B***, Profiles of the G173 daylight standard (white circle) and its 2- and 3-wavelength mimics (black and green arrowheads, respectively), projected onto the 3K map. By changing the intensity ratio of the two G173 mimic wavelengths, different regions of solar elevation can be replicated (blue dots; note that the y axis accounts for only 1.69% of the variance). Using three wavelengths increases the precision of mimicry (pink dots). ***C***, Profiles of artificial light (red circles, closed for measurements and open for CIE standards) projected onto the 3K map. Also shown are G173 profiles with different prereceptoral filtering: adding a cage (yellow circle) and removing the eye’s optics (blue circle). Inset: profiles of monochromatic lights on the periphery of the 3K profiles (gray dots inside the dashed box). For each arrow/arrowhead, the apex point indicates the position of the mimic.

To consider how artificial lights compare to a range of natural skylights, we projected their photopigment stimulation profiles onto the 3K profile map. Most broadband, artificial profiles were far from or on the periphery of the natural profile cloud; xenon was an exception (**Fig. 4*C***, and **Materials and Methods**). The profiles of monochromatic light were distant from the cloud, delineating a broad arc around it (**Fig. 4*C***). Therefore, even when considering the many varieties of ambient solar illumination, most artificial lights are outliers in their actions on the mouse visual system, as judged by distance in a space defined by the stimulation profiles of relevant photopigments (see below for cluster analysis).

We also investigated how artificial lights stimulate mouse photopigments under two conditions that are common in research environments. The first incorporates additional prereceptoral filtering by a “clear” cage (Dauchy et al., 2013). The cage-filtered G173 and xenon profiles fell outside the 3K cloud. We also considered a condition of less prereceptoral filtering: When the retina has been dissected free of the eye, as for *ex vivo* physiological study. The less-filtered xenon and G173 profiles fell outside the 3K cloud or shifted within it, respectively (**Fig. 4*C***).

These analyses underscore the importance of prereceptoral filtering even for natural light. Compensating for prereceptoral filtering of broadband illumination is impractical due to the complexity of the filter and the light. Compensating for monochromatic light is simple, requiring only an adjustment of intensity, but monochromatic light remains unnatural in its effect on the mouse photopigments. On the other hand, the ratios of minimal, few-wavelength mimics can be adjusted for prereceptoral filters of any shape, if the wavelengths are transmitted sufficiently.

### Mimicking and mapping natural light for other dichromatic mammals

We investigated the construction of mimics and maps for other dichromatic mammalian species: rats, cats, rabbits, and dogs. We used published estimates of ocular transmission spectra and λ_max_ values of rod and cone photopigments (Gorgels and van Norren, 1992; Merriam et al., 2000; Douglas and Jeffery, 2014)(**Table S1**). Melanopsin’s individual states have not been defined for these species. However, we have found that the R and E states have similar λ_max_ values in mouse and macaque monkey, suggesting broad evolutionary conservation (Emanuel and Do, 2015; Liu et al., 2023). Furthermore, melanopsin’s λ_max_ appears similar across many mammalian species, from humans to whales (McDowell et al., 2025); as estimated, these λ_max_ values are likely to represent activation from both R and E states, suggesting that their individual λ_max_ values are conserved (Emanuel and Do, 2023). For these reasons, we adopt the λ_max_ values reported for mouse melanopsin’s R, M, and E states (Matsuyama et al., 2012; Emanuel and Do, 2015; Emanuel and Do, 2023).

We tested the possibility of accurately mimicking natural light and its variations for these species using a minimal spectrum. We found that two wavelengths in the correct proportion mimicked the G173 profile with deviation of <5% across all species (RRMSE; **Fig. 5**). For each species except dog, the mimic profile was more similar to the G173 profile than the artificial broadband or monochromatic profiles (**Fig. S2** and **Table S2**). Finally, for all species, adjusting the intensity ratio of the two wavelengths (optimized for the 3K profiles) sufficed to replicate variations of skylight in the 3K cloud (**Fig. S3**) and to correct for different prereceptoral filters (not shown).

**Figure 5.**
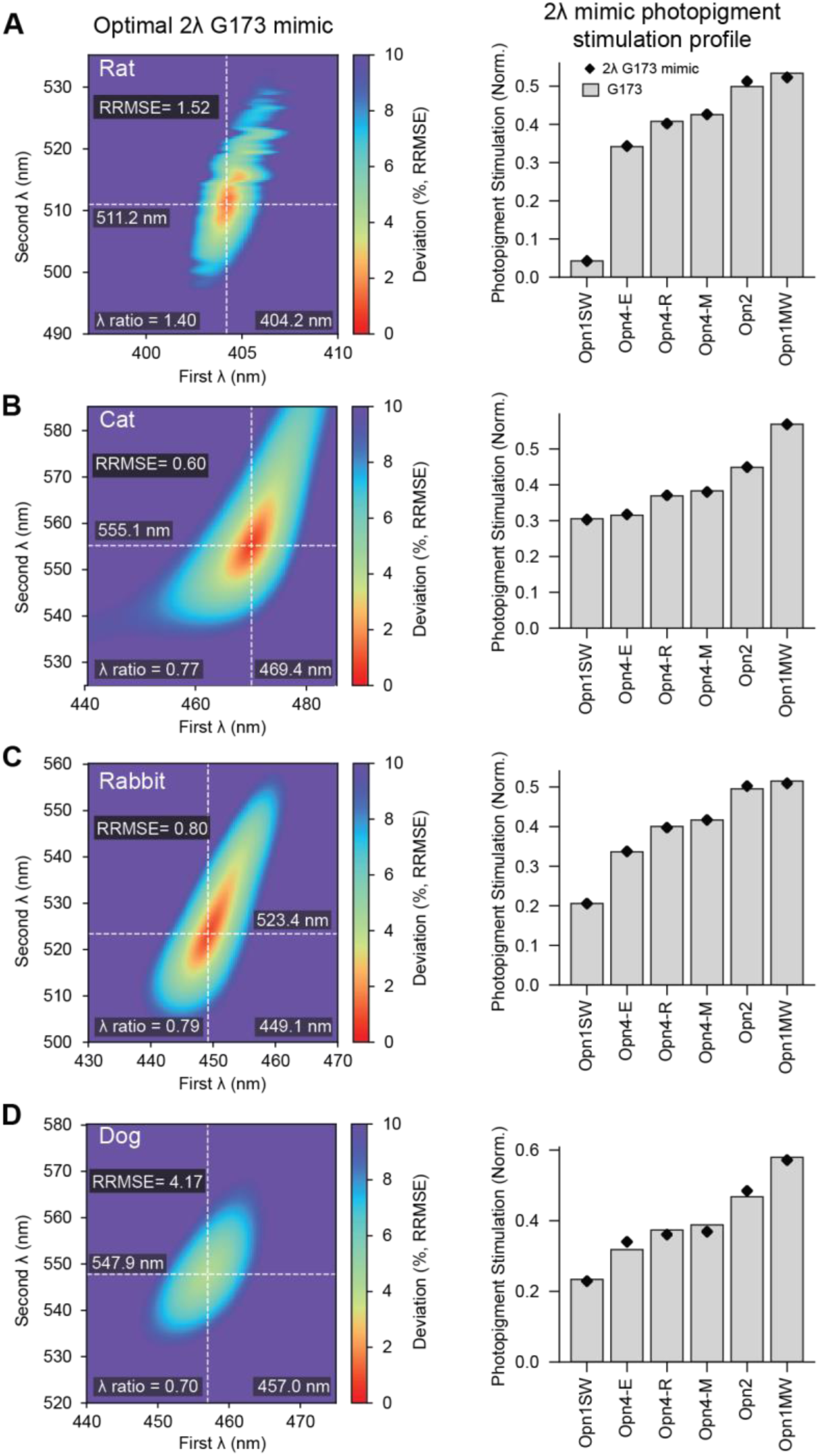
2-wavelength mimicry of natural light for dichromatic mammals (rat, cat, rabbit, and dog). Shown are mimics of the G173 photopigment stimulation profile for rat ***A***, cat ***B***, rabbit ***C***, and dog ***D***. *Left panels:* Effect of varying the two wavelengths on mimic accuracy (their ratios are fixed to optimal values). *Right panels:* The photopigment stimulation profiles of G173 (gray bars) compared to those of the optimal 2-wavelength mimics (black diamonds).

Thus, a 2-wavelength natural-light mimic performs well for the dichromatic mammals tested here. As for the mouse, a third wavelength increased accuracy, though two wavelengths appear sufficient (**Figs. S4*B-E*** and **S5*D***).

Using the 3K natural light spectra (**Fig. 2*A***), relevant ocular transmission spectra and photopigment λ_max_ values, we constructed maps of natural photopigment stimulation profiles for these species (Gorgels and van Norren, 1992; Merriam et al., 2000; Hernandez-Andres et al., 2001; Douglas and Jeffery, 2014; Spitschan et al., 2016)(**Fig. 6**). We found that these maps resembled one another and that of the mouse. For each species, two PCs captured >99.8% of the variance, 3K profiles showed orderly variation across the maps with solar elevation (not shown), profiles for most of our measured, broadband lights bordered or were outside the 3K cloud, and profiles of monochromatic lights surrounded the 3K-profile cloud at a distance (**Fig. 6**). Thus, these maps are largely conserved across dichromatic species despite their varied ocular transmission spectra and photopigment λ_max_ values. A trivial explanation for this conservation is that we used the same λ_max_ values for melanopsin, which accounts for three of the six photopigment states. However, conservation remains when omitting melanopsin (not shown; **Materials and Methods**).

**Figure 6.**
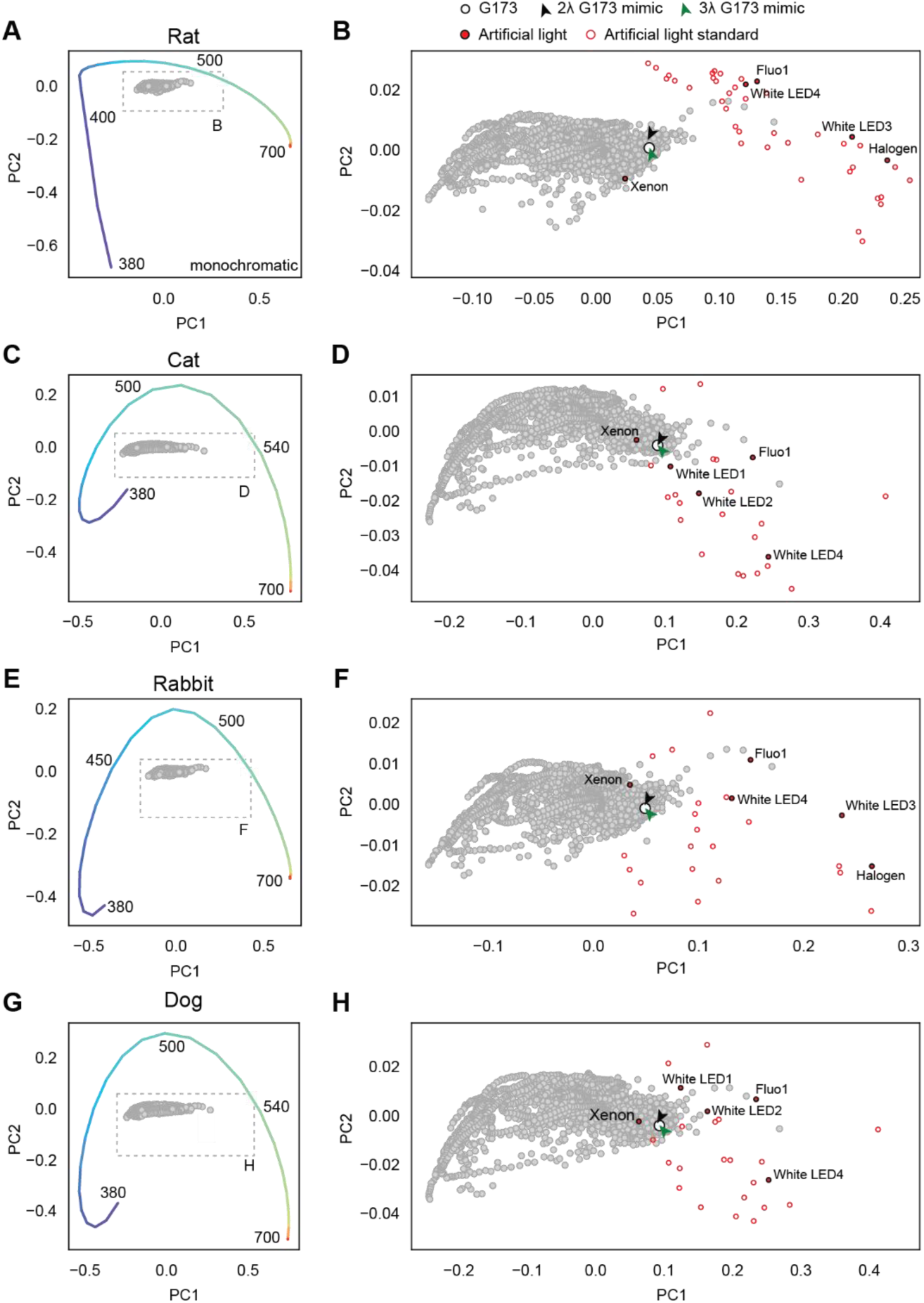
Mapping photopigment stimulation profiles under natural and artificial lights for dichromatic mammals. *Left:* Maps of photopigment stimulation profiles (from the 3K set of natural spectra), with monochromatic lights projected (380–700 nm). *Right:* The boxed region on the left, expanded to show profiles of the 3K spectra (gray dots), the G173 standard (white circle), artificial lights that fall nearby (measured, closed red circles; CIE standards, open circles); and 2- and 3-wavelength mimics (black and green arrowheads). These analyses are shown for rat (***A*** and ***B***), cat (***C*** and ***D***), rabbit (***E*** and ***F***), and dog (***G*** and ***H***).

### Mimicking and mapping natural light for humans

Most humans are trichromatic. We asked whether a small set of wavelengths could reproduce skylight’s stimulation of their seven photopigment states (compared to the six of dichromatic mammals). We used a standard transmission spectrum of the eye (CIE, 2006), and published λ_max_ values for the rod and cone photopigments (**Table S1**; **Materials and Methods**)(Bowmaker and Dartnall, 1980; Merbs and Nathans, 1992). Two variants of the Opn1LW photopigment are common in the population, and we used their average λ_max_ (554.55 nm)(Merbs and Nathans, 1992). For melanopsin, we adopted the λ_max_ values for macaque melanopsin’s R and E states, which appear slightly shorter than those of the mouse (Liu et al., 2023). The M state’s λ_max_ is undefined, so we used that of mouse melanopsin (Matsuyama et al., 2012). The photoswitching of macaque melanopsin is similar to that of mouse melanopsin (Emanuel and Do, 2015; Emanuel and Do, 2023; Liu et al., 2023); because this property depends on the M state’s λ_max,_ any difference in this value between macaque and mouse is likely to be small. We accounted for self-screening of rod and cone photopigments (**Materials and Methods**)(Alpern et al., 1987). We omitted macular pigment to focus on the majority of the retina (but see below).

For a 2-wavelength mimic of G173, we found that the optimum was 470.2 and 557.7 nm in a ratio of 0.64:1. Its profile deviated from that of G173 by <8% for any given photopigment; across photopigments, the RRMSE was 5.53% (**Fig. S5*A*** and ***C***). We also considered a common standard for human daylight perception, CIE D65 (CIE, 2018b). It is a mathematical model with a correlated color temperature of ∼6500 K. Its optimum 2-wavelength mimic comprised 469.3 and 556.0 nm in a ratio of 0.71:1. The profile of the D65 mimic deviated from that of D65 itself by <8% for any given photopigment and 5.20% when averaged across all photopigments (not shown).

Next, we tested mimicry with three wavelengths. For G173 and D65, the optimal wavelengths were 461.7, 523.3, and 592.6 nm in a 0.64:0.77:1 ratio, and 460.9, 522.4, and 591.4 nm in a 0.75:0.85:1 ratio, respectively. These 3-wavelength mimics deviated from G173 or D65 by <0.3% for any photopigment; the average deviations were 0.13 and 0.12%, respectively (**Fig. S5*B-C*,** not shown for D65). Given this high accuracy, we asked if a fixed set of three wavelengths could mimic any of the 3K natural light profiles (**Materials and Methods**). We found that a combination of 460.2, 520.7 and 591.0 nm, with the relative intensity of any one wavelength ranging from 0.21-2.27 (normalized to the last wavelength), was optimal. Across all 3K profiles, this 3-wavelength mimic (with varying ratio) had a photopigment stimulation profile that deviated <1.83% for any photopigment and 0.36 ± 0.16% on average (**Fig. S4*F***). Thus, a 3-wavelength mimic can replicate skylight’s meteorological and daily variations.

We constructed a map of natural photopigment stimulation profiles for humans, using the set of 3K natural light spectra. It differed in shape from those of dichromatic species. Nevertheless, two PCs captured 99.9% of the variance (**Fig. 7*A***). Moreover, 3K profiles showed orderly variation across the maps with solar elevation (not shown); profiles for most artificial, broadband lights bordered or lay outside the cloud; and profiles for monochromatic lights surrounded the cloud at a distance. Hence, most artificial lights, including many that are designed to appear natural to humans, diverge from natural skylight in their stimulation of the retinal photopigments (see below for additional analysis). By contrast, the profiles of the 2- and 3-wavelength mimics of G173 and D65 spectra were near or within the cloud of 3K profiles, with relatively low deviation (RRMSE values; **Figs. 7*A*** and **S5*F***).

**Figure 7.**
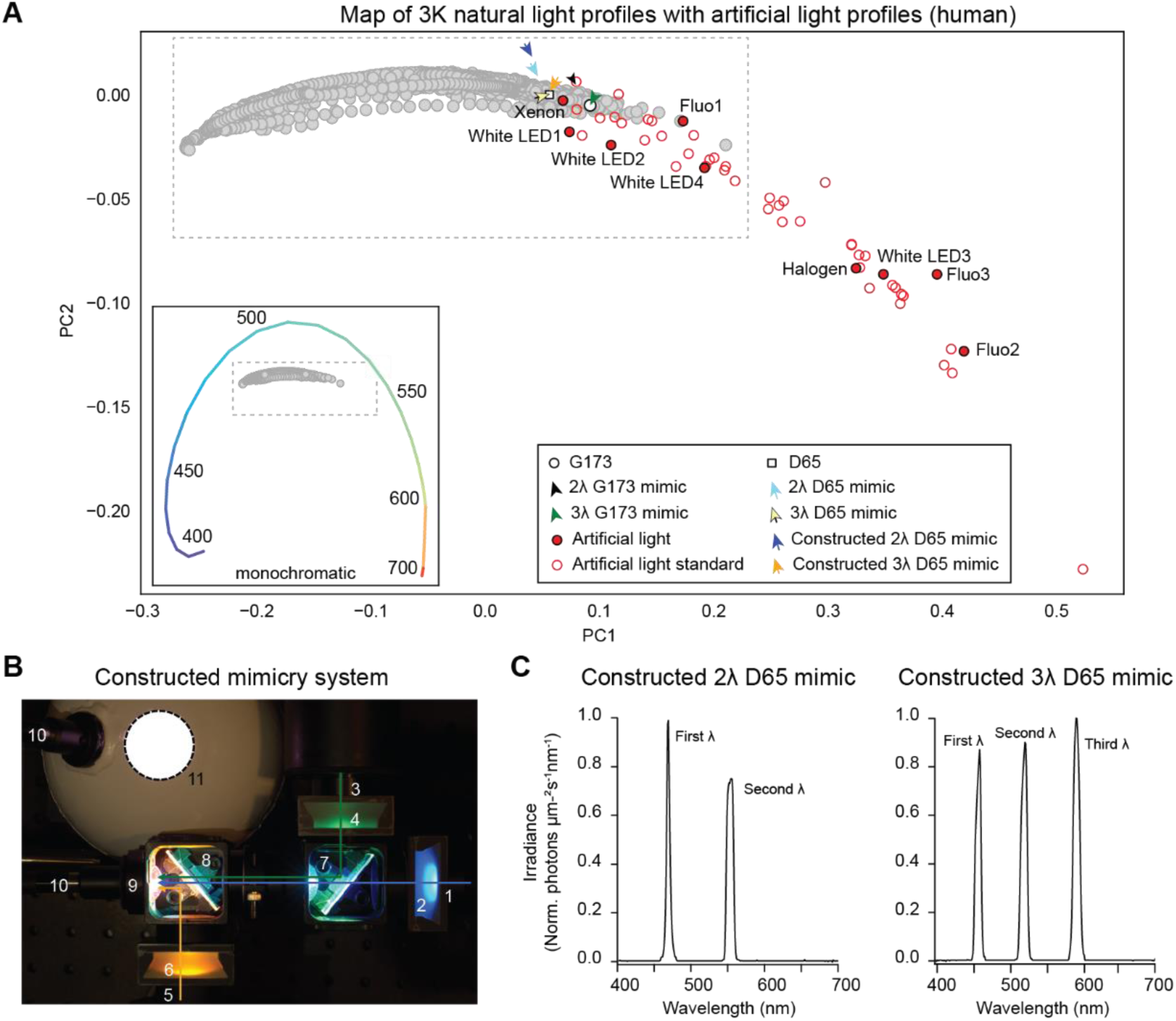
**Mimics and maps of natural light for trichromatic humans. *A***, The map of 3K natural light profiles for humans, shown with profiles of the G173 daylight standard (white circle), 2- and 3-wavelength mimics of G173 (black and green arrowheads), D65 standard (yellow square), optimal 2- and 3-wavelength mimics of D65 (light blue and yellow arrows), constructed 2- and 3-wavelength mimics of D65 (blue and orange arrows), measured and standard artificial lights (closed and open red circles), and monochromatic lights (inset, with the 3K profiles shown as gray dots inside the dashed box; wavelengths shorter than 400 nm are scarcely transmitted by the human eye). For each arrow/arrowhead, the apex point indicates the position of the mimic. See **Table S3** for additional information. ***B***, A constructed mimic of the D65 daylight spectrum, shown in its 3-wavelength configuration with the 12.7-cm Ganzfeld diffusing sphere. Labeled are three narrowband light sources (**1**, **3**, and **5**), bandpass filters (**2**, **4**, and **6**), beamsplitters (**7**, **8**), a lens (**9**), a light guide (**10**), and the Ganzfeld sphere (**11**; aperture is outlined). ***C***, Measured output spectra of the 2- and 3-wavelength mimic systems.

### Comparison of natural and artificial photopigment stimulation profiles

To assess differences between natural and artificial profiles that may lie outside the two dimensions of the 3K maps, we applied cluster analysis to all dimensions; six for dichromatic species and seven for humans (**Materials and Methods**). We detail the outcomes in **Table S3**. For each species, >99% of the 3K profiles were classified as part of one cluster, along with the G173 standard. Across all species, profiles of most artificial lights were excluded from this core: >95.2% for standard spectra and all except xenon for our set of measured spectra. Furthermore, when G173 was filtered by cage material or lacked filtering by the eye (see above), its profile was often excluded from the core. This analysis underscores the abnormal effect of most direct artificial light on the mammalian photopigments, and the importance of prereceptoral filtering on skylight.

The core 3K profiles clustered with profiles of all 3-wavelength mimics for all species and across all conditions assessed (i.e., *in vivo*, *ex vivo*, and cage) and the 2-wavelength mimics of mouse and cat (for all conditions assessed; **Table S3**). The profiles of the remaining mimics deviated little from those of natural light (e.g., <5% RRMSE for G173). Their exclusion likely reflects the clustering method’s sensitivity to covariation; photopigment stimulations covary across natural spectra in a way that can be discontinuous with the profile induced by the mimic. This discontinuity is subtle compared to the qualitatively different profiles under most of the common sources of artificial light.

### Construction and evaluation of human natural light mimics

We designed, built, and tested 2- and 3-wavelength skylight mimics for humans. Actual illumination spectra are broader than the theoretical, single wavelengths given above. Therefore, we asked how deviations from theoretical wavelength and ratio optima affected mimic accuracy (as illustrated for the mouse in **Fig. 1*C-D***). We found regions of shallowly changing, relatively low average deviation (RRMSE <10%) and designed our mimics around them (**Materials and Methods**). We selected two or three LEDs, collimated their outputs, sent the beams through bandpass filters, combined the beams with dichroic mirrors, focused the combined beam onto one end of a liquid light guide, and connected the opposite end to a custom Ganzfeld sphere (**Fig. 7*B***; **Materials and Methods**).

We used a Ganzfeld because it produces diffuse illumination, like ambient natural light. This stimulus is also precisely quantifiable, as each point in the retina receives an intensity that is nearly the product of the Ganzfeld’s output, the attenuation by the eye’s optics (cornea, iris, and lens), and the inverse of the retinal surface area. Our Ganzfelds had a 30.5- or 12.7-cm outer diameter, a thick internal shell of BaSO_4_ (which reflects relevant wavelengths evenly), and a 13-or 3.4-cm diameter viewing port (the former filling much of one’s field of view and the latter achieving higher intensities; **Materials and Methods**). We measured light at the viewing port of the Ganzfelds with a calibrated spectrometer and power meter, converting units to photons µm^-2^ s^-1^ nm^-1^, and used those measurements for evaluation.

We focused on the D65 daylight spectrum because it is a common standard for human vision (Lucas et al., 2024). We found that the constructed mimics performed close to our theoretical predictions. For the constructed, 2-wavelength D65 mimic, the outputted spectra had peak wavelengths at 467.7 nm (full-width at half-maximum, FWHM, of 4.7 nm) and 554.5 nm (10.5 nm) in a ratio of 0.69:1 (integrals of the photon flux density across wavelength; not shown).

Compared to the D65 standard, this constructed mimic had a maximum deviation of <8.5% for any given photopigment and an average deviation of 5.35% (compared to 5.20% for the theoretical 2-wavelength D65 mimic, see earlier). For the constructed, 3-wavelength D65 mimic, the peak wavelengths were 460.0 nm (FWHM of 8.6 nm), 522.6 nm (9.6 nm), and 592.6 nm (9.8 nm) (**Fig. 7*C***). Their ratios were 0.72:0.92:1. This mimic had a maximum deviation of <1.6% for any given photopigment and an average deviation of 0.86% (compared to 0.12% for the theoretical 3-wavelength D65 mimic, see earlier). The constructed 2- and 3-wavelength mimics had profiles that fell near or within the cloud of 3K natural light profiles, respectively (**Fig. 7*A***), and the latter clustered with the core.

The smaller Ganzfeld gave a maximum output of 6.5×10^7^ photons µm^-2^ s^-1^ (integrated across the measured spectrum). Using the CIE photopic luminosity function (CIE, 2019a), this corresponds to ∼11,000 lux and is equivalent to ambient illumination at ∼10-20° of solar elevation (Cronin et al., 2014). The maximum output of the 30.5-cm Ganzfeld was ∼1,100 lux (ca. 0° solar elevation). Both outputs meet safety standards (IES, 2020). They can be controlled finely over a broad range—for example, by using currents of different amplitude, pulse-width modulation, and neutral density filters—and their maximum outputs increased, if desired, using more intense sources.

We designed our natural light mimics for all photopigments that are known to trigger electrical signals in humans, thus replicating light’s actions on both image vision (perception, which is primarily mediated by rod and cone photopigments) and non-image vision (e.g., circadian photoregulation, for which melanopsin is vital)(Lucas et al., 2014; Do, 2019; Lucas et al., 2024). We consider nearly as many photopigment states for the former (four) as the latter (three). How does optimizing for all photopigments influence visual perception? We addressed this question by tuning our D65 mimics for the central 10° of visual field, akin to the standard observer commonly used to evaluate color space (CIE, 2006). Macular pigment is substantial in this field so we considered its prereceptoral filtering (**Materials and Methods**)(CIE, 2006). We kept the wavelengths constant and adjusted their ratios. We found that the optimal 2- and 3-wavelength mimics had ratios of 0.72:1 and 0.73:0.87:1, respectively. We tested the outputs of the constructed mimics and found that the 2-wavelength version deviated by <8.9% for any photopigment and 5.75% across photopigments (RRMSE), while the 3-wavelength version deviated by <1.29% for any photopigment and 0.70% across photopigments. Next, we mapped the outputs of these mimics to color space (**Fig. 8*A-B*** and **Materials and Methods**)(CIE, 2018b). The distances between D65 and the 2- and 3-wavelength mimics were 25.69 and 2.46 (CIEDE2000). The latter is considered barely perceivable. Collectively, these analyses indicate that three wavelengths mimic skylight well for both image and non-image vision (**Fig. 8*C***).

**Figure 8.**
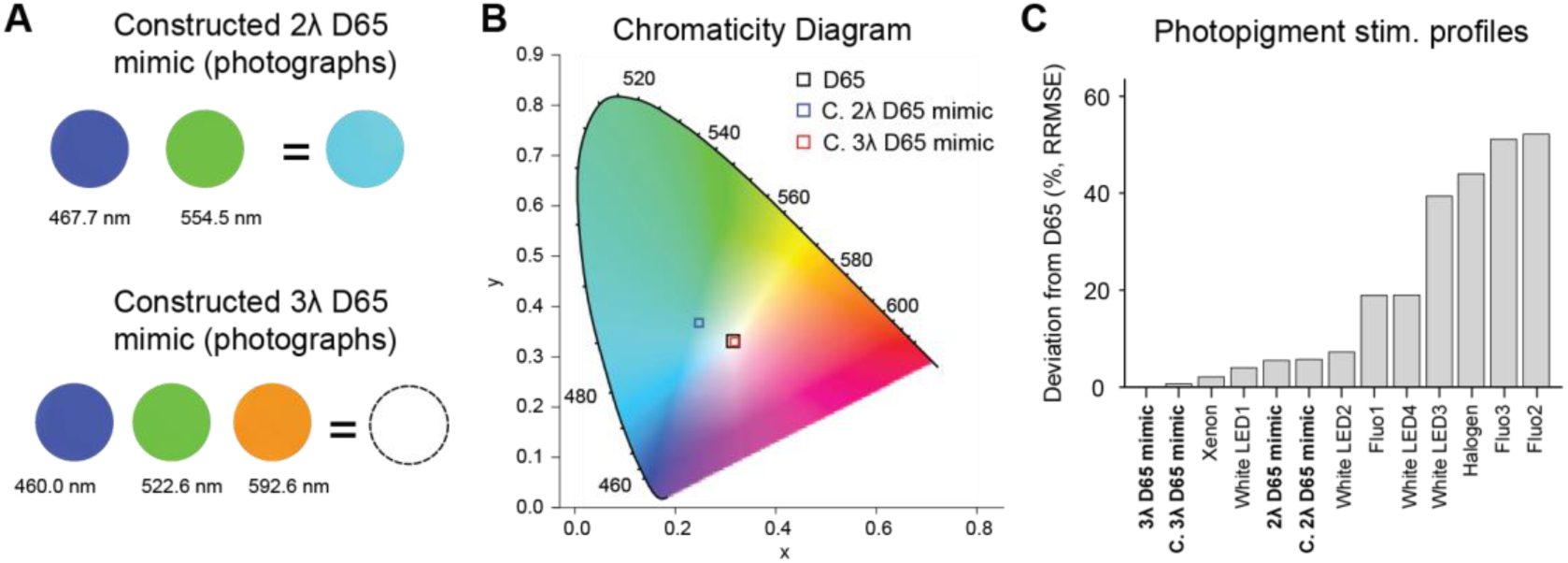
**Mimics of natural light in human color space. *A***, Photographs of the Ganzfeld sphere’s aperture (see Fig. 7B*) showing* the individual light sources and their combination, for the 2- and 3-wavelength mimics. ***B***, Projection of D65 and its constructed 2- and 3-wavelength mimics (2λ D65 mimic and 3λ D65 mimic in Fig. 7C) onto the CIE chromaticity diagram (a map of human color perception)(CIE, 2019b)**. *C***, Comparison of the D65 photopigment stimulation profile with those of measured artificial lights. Deviation is expressed as the relative root mean square error, RRMSE, across photopigments. Also see **Fig. S5*F*.**

## DISCUSSION

We designed and built artificial light sources that mimic photopigment stimulation profiles produced by natural skylight. These sources perform well with only two wavelengths and almost perfectly with three, and are readily adjusted to mimic variations of natural light and to compensate for prereceptoral filtering. We have also examined how thousands of natural light variations are represented by the photopigment complements possessed by humans, mice, and other mammals. We report that the representation for each species is captured in a 2-dimensional map that has orderly and informative substructure. Here, we place our work into context and discuss how it may be extended.

### Mimicking natural light’s stimulation of mammalian photopigments

By accounting for how an animal’s complement of photopigments is stimulated by light, one can manipulate light to produce desired stimulation patterns. This is the basis of silent substitution, in which light’s spectrum and intensity are varied to selectively modulate one or more photopigments while keeping the others in a constant state (Spitschan et al., 2017b; Mouland et al., 2019). Silent substitution has provided insight into the roles of specific photopigments and their interactions, including summation and opponency, even without manipulations like gene knockouts that are unfeasible in humans and other species (Dacey et al., 2005; Spitschan et al., 2014).

We have used similar principles to develop artificial lights that mimic natural light’s effects on the mammalian photopigments. Different spectra that produce the same percept are called metamers. Our mimics are metamers for perception as well as for non-perceptual actions of light. They differ in several respects from prior work in a similar vein (Mouland et al., 2019): we mimicked natural light for species from mice to humans and considered prereceptoral filtering for all; incorporated melanopsin’s three stable conformations; included many natural light variations; developed spectra whose wavelength bands are narrow and few in number; identified a simple means of correcting for prereceptoral filtering; omitted potentially harmful, ultraviolet wavelengths; and verified human natural light mimicry using psychophysical color space. The natural light mimics that we describe are accurate, safe, and flexible. We anticipate that they can be crafted for many species.

These mimics are of natural skylight that is weighted by the spectral sensitivities of relevant photopigment states. Other factors should be engaged naturalistically within this context, including each photopigment’s extinction coefficient, quantum efficiency of isomerization, expression level, expression locus, and biological actions. The mimic is also agnostic to the presence of a photopigment or photopigment state for which it was designed. For example, only one melanopsin state is found in darkness; formation of the other states depends on the spectrum, intensity, and timing of illumination; and the pigment bleaches (Walker et al., 2008; Do et al., 2009; Emanuel and Do, 2015; Tsukamoto et al., 2015; Liu et al., 2023). The mimic acts on photopigment states when they are available. Furthermore, one may control the mimic’s spatial and temporal properties, and the photopigments should respond accordingly. Spatial control of mimicry is also important when prereceptoral filtering varies with retinal location, due to anisotropic features like pigmentation and vasculature (Snodderly et al., 1984a; Snodderly et al., 1984b; Spitschan et al., 2015).

An advantage of mimicking natural light with only two or three wavelengths is the simplicity of correcting for prereceptoral filtering. A disadvantage is that a minimal mimic’s reflected light is, in many contexts, unnatural. Minimal mimics of the kind we introduce here are most effective when delivered directly, such as for light therapy and research; when the intensity of direct illumination outstrips that of reflected light; or when light is altered little after reflection (as in a white room). Achieving natural reflectance spectra is likely to require illumination with broader spectra. One might find a middle ground depending on the application. For example, many LEDs that appear white and produce naturalistic reflectance spectra have one or more broad peaks. Broadening one peak of our human D65 mimic increases its accuracy (from an average deviation across photopigments of 5.75 to 3.1%; not shown).

### Incorporating additional photopigments into mimics of natural light

The natural light mimics we present are comprehensive in that they include melanopsin’s three states, but they omit molecules that are not known to trigger electrical signals. A notable example is neuropsin (Opn5), which has two stable states (λ_max_ of 380 and 470 nm), is found in the mammalian retina and brain, and is implicated in processes that include photoregulation of the retina clock and thermogenesis (Calligaro et al., 2022). We considered how minimal natural light mimics perform when adding neuropsin’s two states, which brings the total number of relevant absorption spectra to eight for mice and nine for humans. We found that 2- and 3-wavelength mimics of G173 have average deviations of 11.11 and 0.69% for mice, and of 14.3 and 1.9% for humans, respectively. Therefore, minimal mimics appear possible even for large numbers of absorption spectra.

### Representing light in terms of photopigment stimulation

Expressing light in terms of its photopigment effects is a basic practice in vision research. For example, light intensity is often given as photoactivated rhodopsin molecules (R*) per rod. One obtains this value by using the rod’s effective collecting area (in units of µm^2^), which incorporates light’s spectrum (in photons µm^-2^ s^-1^ nm^-1^) and prereceptoral filtering, as well as rhodopsin’s orientation, extinction coefficient, quantum efficiency of isomerization, and density (Luo et al., 2008). The resulting units of R*/rod focuses on the level of biological activation rather than properties of the light itself. This measure is possible because rods are stereotyped and have been characterized extensively across species; it is also available for some cones. A measure that requires fewer parameters is the α-opic irradiance, obtained by integrating the product of the absorption spectrum of each photopigment (α) of a species (corrected for prereceptoral filtering) and the power spectral density of the light (watts per unit area and unit time) at the retina. For intuition, an additional transformation yields the intensity of standard daylight (D65) that would give the same photopigment-weighted irradiance (α-opic equivalent daylight illuminance; EDI)(Lucas et al., 2024).

Our work extends these traditions. Like R*/rod, we use photons rather than watts to focus on biological effects. Like α-opic irradiance, we consider several photopigment types across species. We go further by incorporating melanopsin’s three stable and spectrally distinct conformations, which were defined relatively recently and have stark effects on melanopsin-expressing cells (Matsuyama et al., 2012; Emanuel and Do, 2015; Emanuel and Do, 2023; Liu et al., 2023). Like α-opic EDI, we examine natural light. However, rather than examining an idealized daylight spectrum like D65, we analyze thousands of spectra that detail the actual, ambient illumination from the sun. In this manner, we produced maps of photopigment stimulation profiles across variations of skylight for mice, rats, cats, dogs, rabbits, and humans. The profile map is likely to be useful. For example, by situating artificial lights within it, one sees how naturalistically they influence perception and regulate physiology.

We expect that our approach can be brought to non-mammalian species, where interesting complications are found. For instance, prereceptoral filtering may be complex; some birds and reptiles have oil droplets of different colors that shape the light received by cones, and other species situate their photopigments deep in the brain or beneath other tissues (Toomey and Corbo, 2017; Calligaro et al., 2022). Some species have many photopigment types—the mantis shrimp has a dozen in its compound eye, and a fruit fly has eight in its compound eye and dorsal ocellus (Thoen et al., 2014; Christenson et al., 2022). Some photopigments are associated with molecules that broaden their spectra, and others change their absorption spectra by switching their chromophore type (Isayama et al., 2006; Corbo, 2021). Examining the photopigment stimulation maps of these species would provide insight into their visual ecologies.

Future mimics and maps might account for ambient solar illumination spectra from other geographical locations, meteorological conditions, and seasons; include spectra of the reflected light that an organism tends to experience; consider the daylight available to aquatic organisms; and broaden this analysis to include other natural illumination such as the spectra of moonlight, starlight, bioluminescence, fluorescence, and chemiluminescence (Johnsen, 2012; Maloney, 1986). These mimics and maps would support investigations of the diversity of photoreceptive mechanisms.

Photopigment stimulation maps reveal how different photic environments influence a given species. It would be interesting and likely useful to determine how the maps of interacting species overlap, and how those overlaps vary with natural conditions as well as with the type and prevalence of artificial illumination.

### Closing remarks

Newton found that passing white daylight through a prism yields a spectrum of colors. Helmholtz discovered that combining complementary colors reproduces the perception of white light (Helmholtz and Southall, 1924). Rod and cone photopigments provided an explanation, but psychophysical experiments suggested that this explanation may be incomplete (Danilova and Mollon, 2022). Behavioral and physiological experiments also alluded to an additional photopigment, which is melanopsin (Do, 2019). We provide evidence that two wavelengths suffice to mimic the effect of natural light even when considering its many variations and melanopsin’s complexities. One advantage of such minimal mimics is easy compensation for prereceptoral filtering, which allows it to be precisely matched across varied conditions—for instance, to study animals and *ex vivo* tissues using the same effective illumination, and to overcome age-related changes in the eye’s optics and hence restore the perception and physiological actions of light from earlier life.

## AUTHOR CONTRIBUTIONS

Conceptualization: 2-wavelength mimics of natural light with compensation for arbitrary prereceptoral filtering, M.T.H.D; maps of photopigment stimulation profiles, P.M.; the remainder: P.M. and M.T.H.D. Experiments and analyses: P.M. Writing: P.M. and M.T.H.D.

## ACKNOWLEDGMENTS

Javier Hernández-Andrés kindly provided metadata of the Granada natural light dataset. Andreas Liu gave an example of wavelength optimization. Franklin Caval-Holme, Sophia Wienbar, and Navid Mousavi provided critical discussion. Grigory Loginov and Navid Mousavi assisted with construction of the diaphragm adaptor for spectrophotometry. Samuel Brill-Weil, Franklin Caval-Holme, Grigory Loginov, Nguyen-Minh Viet, Victoria Amstrup Vold, and Sophia Wienbar provided critiques of the manuscript. This research was funded by NIH grants EY023648, EY025555, EY032731, EY030628, and EY034089 (to M.T.H.D.). P.M. is a Warren Alpert Distinguished Scholar in Neuroscience, a Lefler Fellow of Harvard Medical School, and a Tommy Fuss Fellow of Boston Children’s Hospital.

## CONFLICT OF INTEREST STATEMENT

Philippe Morquette and Michael Tri H. Do declare that a patent application related to the work described in this manuscript has been filed through Boston Children’s Hospital.

## SUPPLEMENTARY MATERIAL

**Supplemental Table 1.** Wavelengths of peak absorption (λ_max_) for the photopigments used in this study. Those of melanopsin states are from the mouse (Matsuyama et al., 2012; Emanuel and Do, 2015), with the exception of the human R and E states, which are taken from the macaque (Liu et al., 2023).

| Species | Photopigment | $\lambda_{\max}$(nm) | References |
| --- | --- | --- | --- |
| **Mouse** | Opn1SW | 358 | Tsutsui et al., 2007 |
|  | Opn4 (E,R,M) | 453.0, 471.0, 476.0 | Emanuel and Do, 2015; Matsuyama et al., 2012 |
|  | Opn2 | 502 | Sakurai et al., 2007 |
|  | Opn1MW | 508 | Sun et al., 1997 |
| **Rat** | Opn1SW | 358 | Yokoyama et al., 1998 |
|  | Mel (E,R,M) | 453.0, 471.0, 476.0 | Emanuel and Do, 2015; Matsuyama et al., 2012 |
|  | Opn2 | 498 | Bridges, 1959 |
|  | Opn1MW | 509 | Radlwimmer and Yokoyama, 1998 |
| **Cat** | Opn1SW | 450 | Guenther & Zrenner, 1993 |
|  | Opn4 (E,R,M) | 453.0, 471.0, 476.0 | Emanuel and Do, 2015; Matsuyama et al., 2012 |
|  | Opn2 | 501 | Guenther & Zrenner, 1993 |
|  | Opn1MW | 553 | Sun et al., 1997 |
| **Rabbit** | Opn1SW | 421 | McDowell et al., 2025 |
|  | Opn4 (E,R,M) | 453.0, 471.0, 476.0 | Emanuel and Do, 2015; Matsuyama et al., 2012 |
|  | Opn2 | 502 | Bridges, 1959 |
|  | Opn1MW | 509 | Radlwimmer and Yokoyama, 1998 |
| **Dog** | Opn1SW | 428 | Jacobs, 1993; McDowell et al., 2025; Neitz et al., 1989 |
|  | Opn4 (E,R,M) | 453.0, 471.0, 476.0 | Emanuel and Do, 2015; Matsuyama et al., 2012 |
|  | Opn2 | 506 | Parkes et al., 1982 |
|  | Opn1MW | 554 | Jacobs, 1993; McDowell et al., 2025; Neitz et al., 1989 |
| **Human** | Opn1SW | 426.3 | Merbs and Nathans, 1992 |
|  | Opn4 (E,R,M) | 453.51, 467.21, 476.02 | Liu et al., 2023; Matsuyama et al., 2012 |
|  | Opn2 | 497.6 | Bowmaker and Dartnall, 1980 |
|  | Opn1MW | 529.7 | Merbs and Nathans, 1992 |
|  | Opn1LW | 552.4 and 556.7 | Merbs and Nathans, 1992 |

**Supplemental Table 2.**
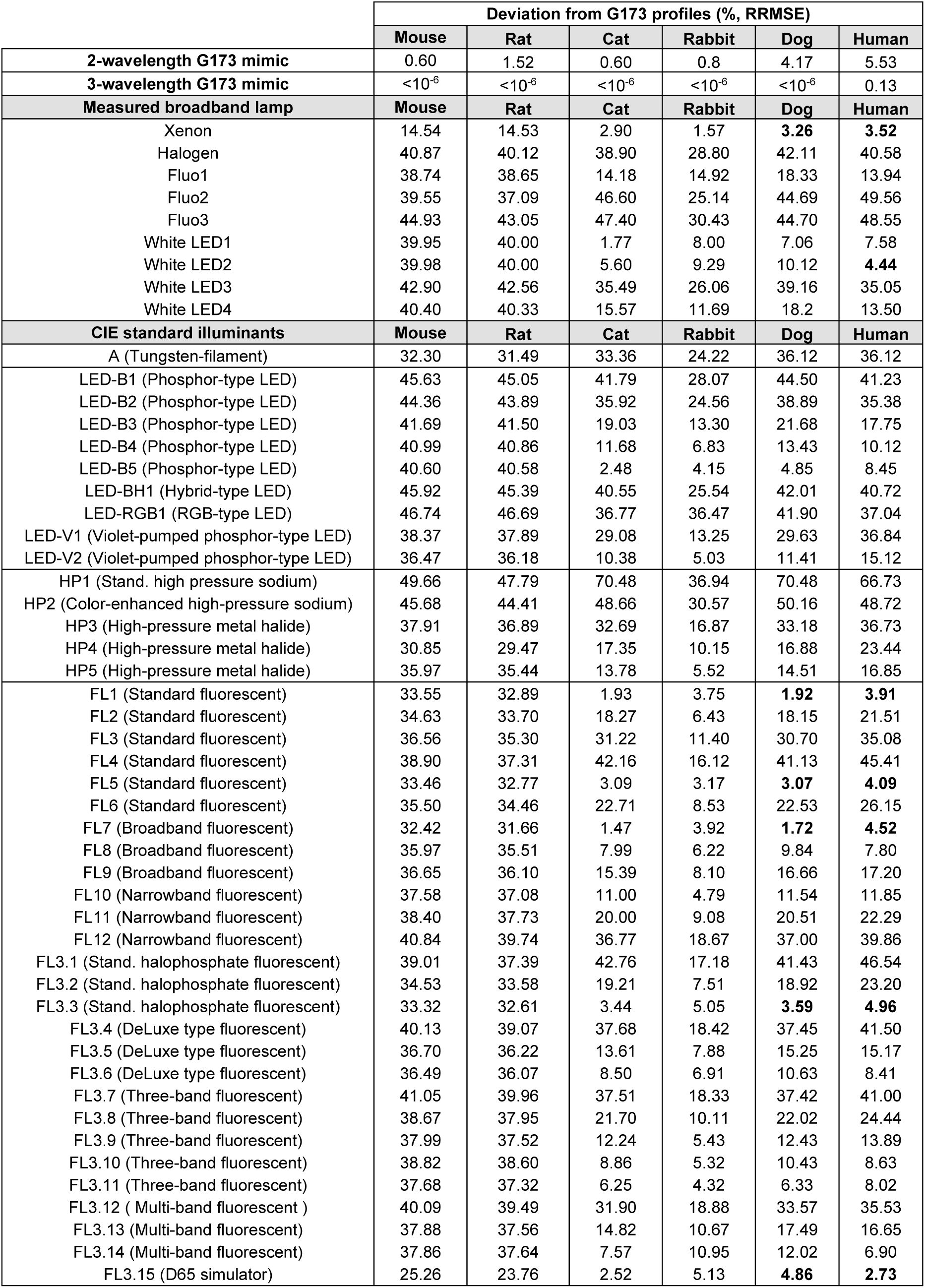
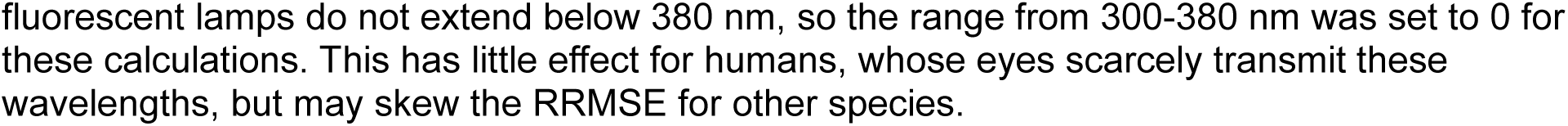
Comparison of photopigment stimulation profiles (%, RRMSE) between G173, 2- and 3-wavelength G173 mimics, measured artificial light sources, and CIE artificial light standards (CIE, 2018b). A range of 300-700 nm was considered. CIE standards for fluorescent lamps do not extend below 380 nm, so the range from 300-380 nm was set to 0 for these calculations. This has little effect for humans, whose eyes scarcely transmit these wavelengths, but may skew the RRMSE for other species.

**Supplemental Table 3.**
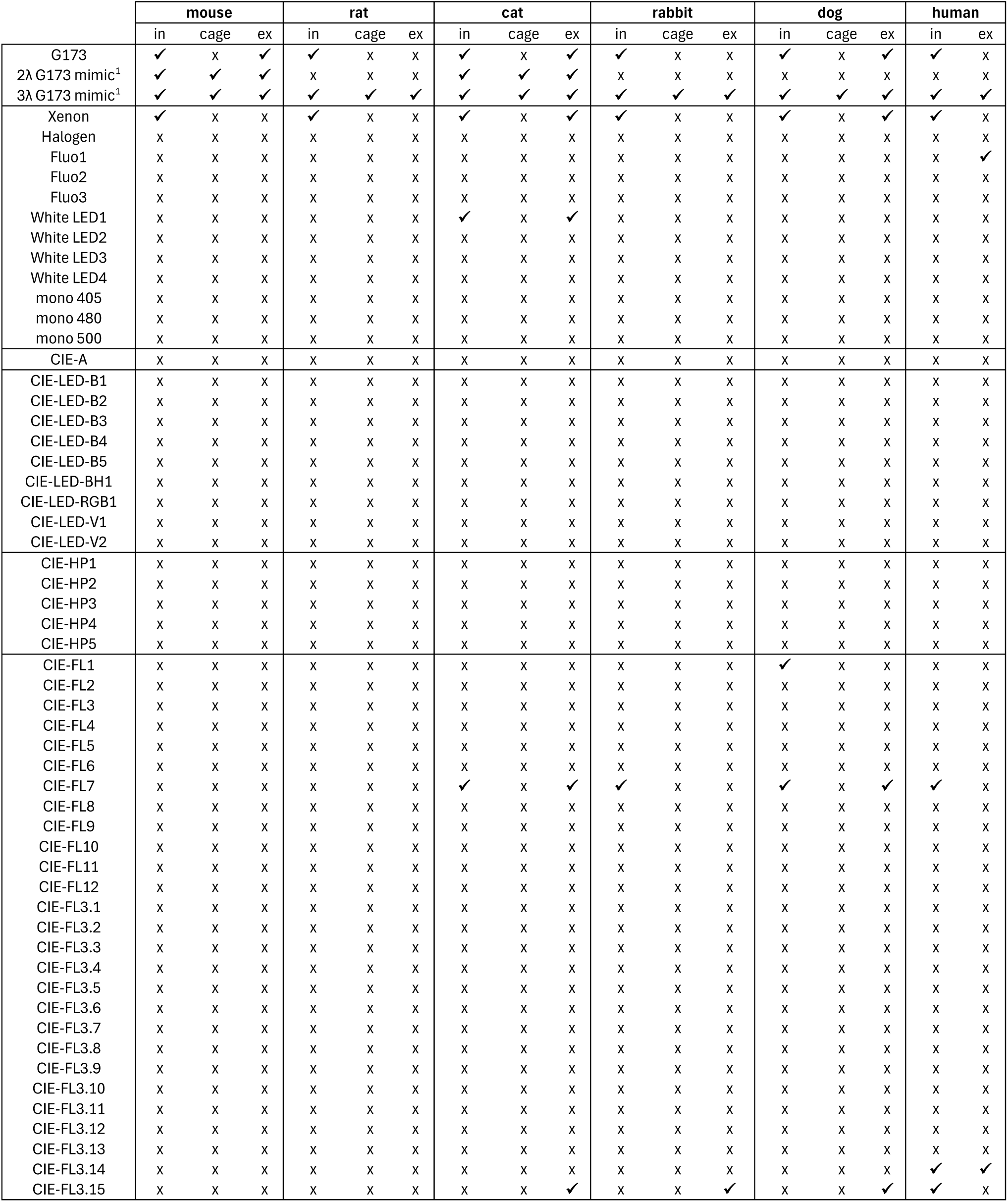
Most photopigment stimulation profiles for artificial light (x) do not cluster with the profiles for natural light (3K set). Some do (✓) but rarely for all conditions (*in vivo*, cage, and *ex vivo*). Profiles for the 2-wavelength (2λ) G173 mimics cluster with natural profiles for all conditions in mouse and cat. The 3λ G173 mimics cluster with natural profiles for all conditions and species. ^1^Note that the mimics have wavelengths and wavelength ratios that are specific to each species and condition; for a given species, the wavelengths were fixed and the ratios changed according to condition.

**Supplemental Figure 1.**
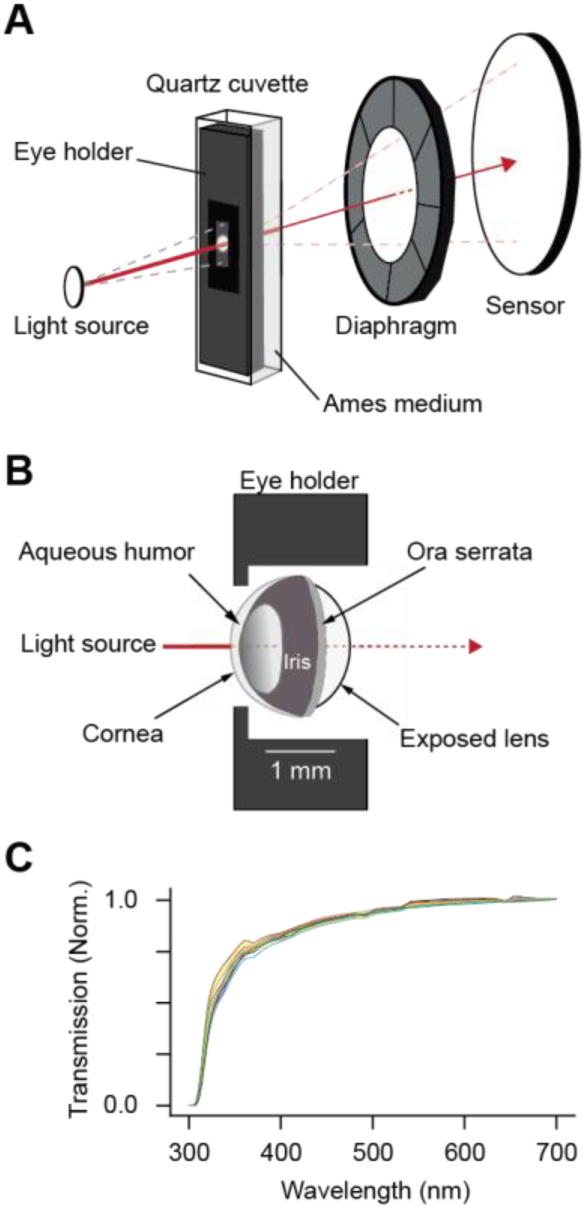
T**r**ansmission **spectrum of the mouse eye. *A***, Schematic of the optical path, inside a spectrophotometer, for measuring the transmission spectrum of the anterior chamber. The anterior chamber was mounted in a custom cuvette filled with oxygenated media, and all transmitted light was captured by the detector. ***B***, Illustration of the anterior chamber’s mounting within the cuvette. ***C***, Raw transmission spectra from 11 adult, wild-type eyes (from 7 males and 4 females), normalized at 700 nm.

**Supplemental Figure 2.**
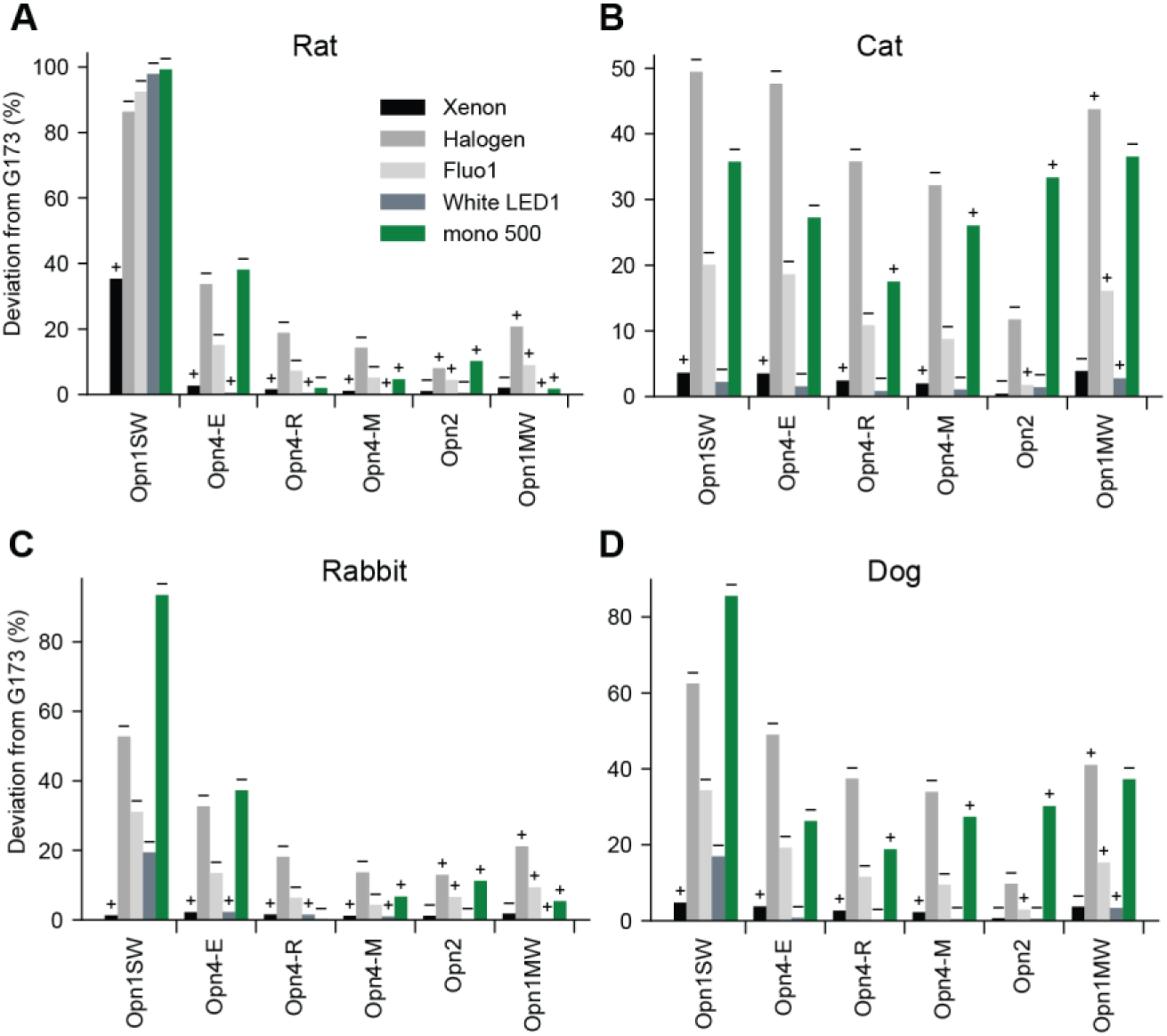
Comparing photopigment stimulations for the G173 daylight standard and example artificial lights for rat (***A***), cat (***B***), rabbit (***C***) and dog (***D***). Artificial lights produced more (+) or less (-) stimulation than G173.

**Supplemental Figure 3.**
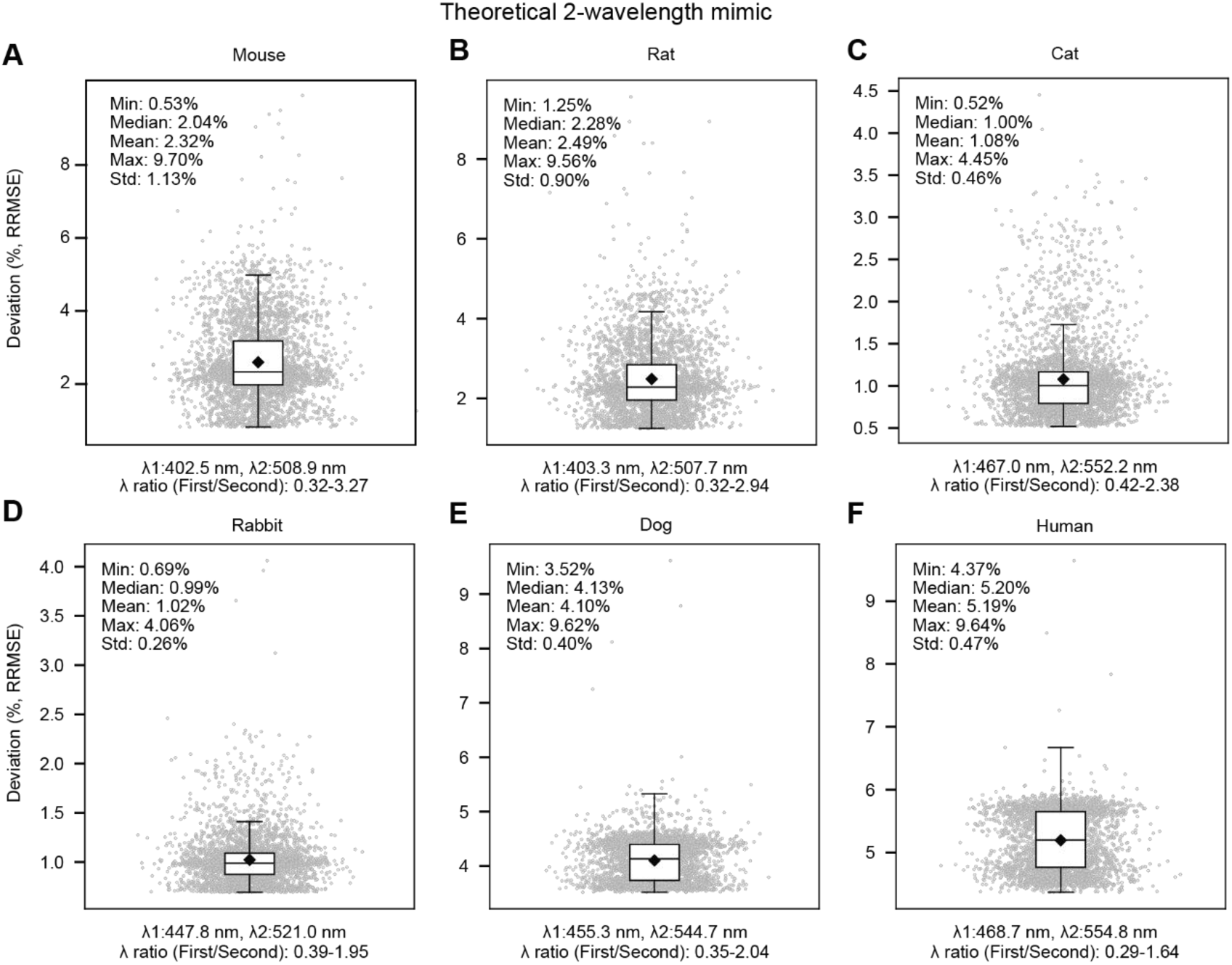
2-wavelength mimicry of natural light’s variations for dichromatic and trichromatic mammals. Two optimal wavelengths, with their intensity ratio adjusted (wavelengths and ratio range are given below each plot), mimic most 3K daylight profiles (gray points) with <10% deviation. Points are spread along the x axis for visualization. Shown are the interquartile range (IQR, box), median (line), mean (diamond), and last point within 1.5×IQR. Analyses are shown for mouse (***A***), rat (***B***), cat (***C***), rabbit (***D***), dog (***E***), and human (***F***).

**Supplemental Figure 4.**
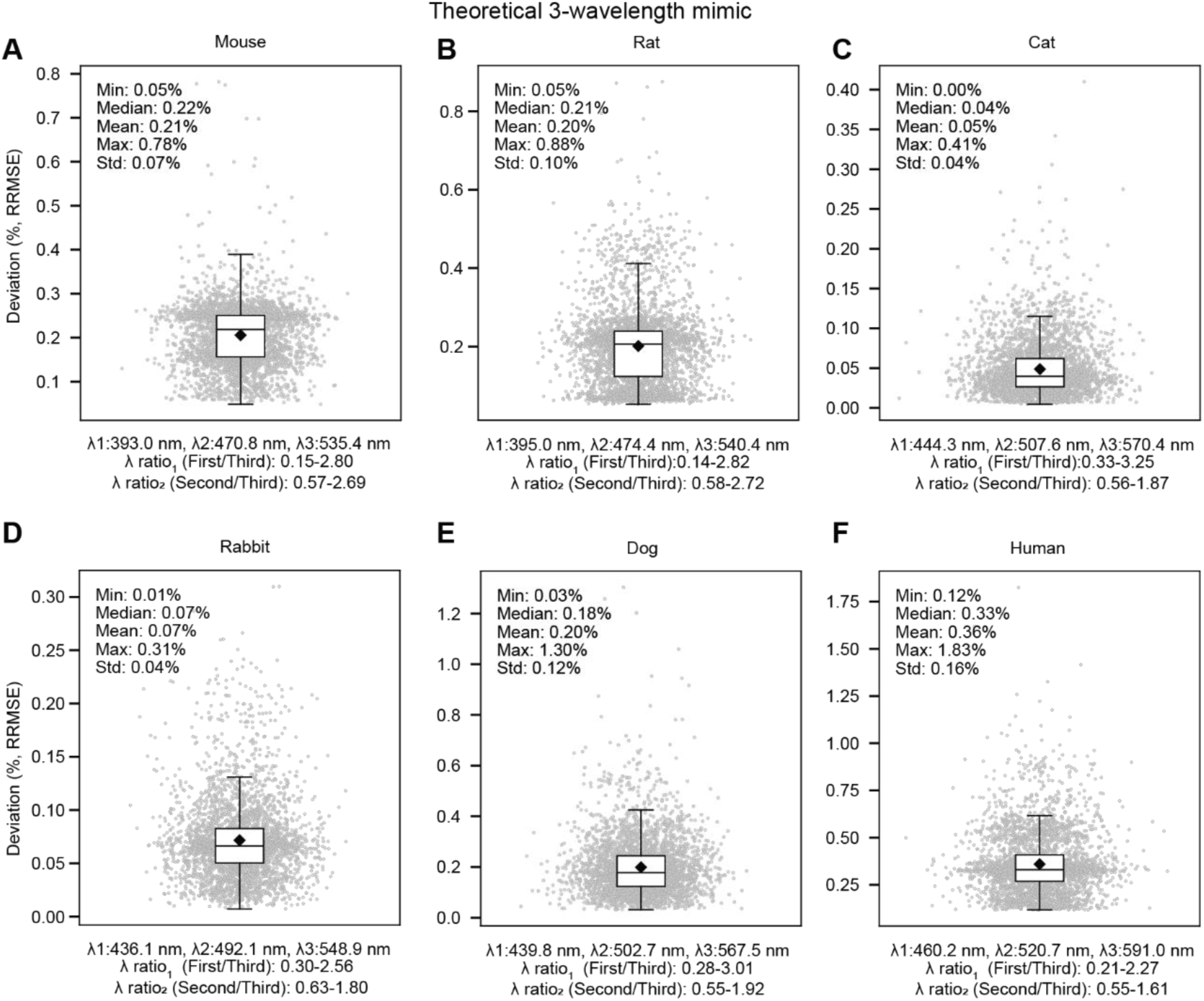
T**h**ree**-wavelength mimicry of natural light’s variations for dichromatic and trichromatic mammals.** As for **Fig. S3**, but for 3-wavelength mimics.

**Supplemental Figure 5.**
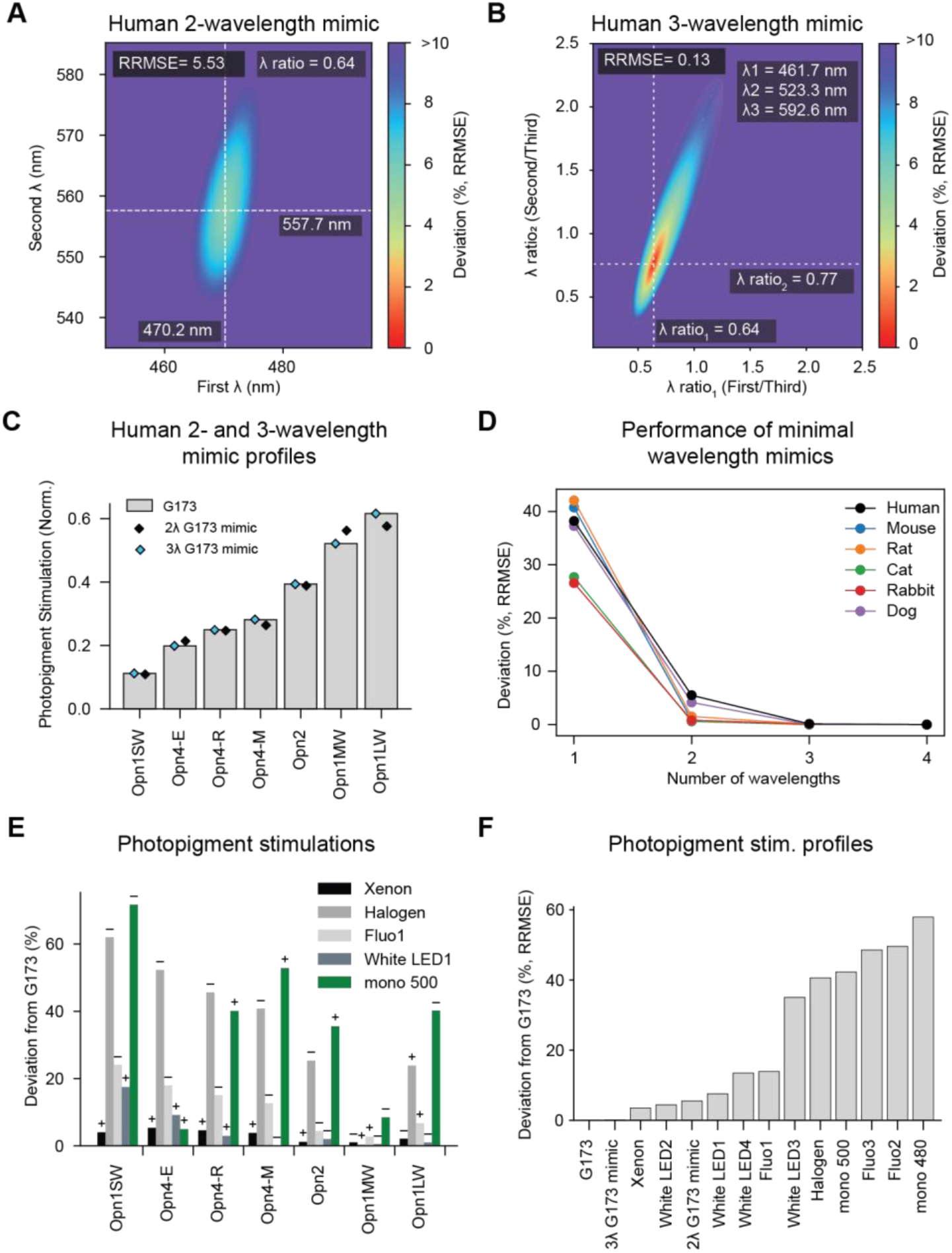
M**i**micking **natural light and assessing artificial light for trichromatic humans. *A***, Effect of varying wavelengths on the accuracy of a 2-wavelength G173 mimic (their ratios are fixed at optimal values). ***B***, Effect of varying ratios on the accuracy of a 3-wavelength G173 mimic (the three wavelengths are fixed at optimal values). ***C***, The photopigment stimulation profile of G173 (gray bars) compared to those of the optimal 2- and 3-wavelength mimics (black and cyan diamonds). ***D***, For all tested species, mimic performance peaks with three wavelengths. ***E***, Artificial lights stimulate human photopigments more (+) or less (–) than the G173 daylight spectrum. ***F***, Deviation of photopigment stimulation profiles of artificial lights from that of G173. The 3-wavelength mimic is more accurate than any tested, artificial spectrum. The 2-wavelength mimic has higher deviation than xenon or one type of white LED, but is preferable to these broadband sources when compensating for different degrees of prereceptoral filtering.

## Notes

### Summary of Updates

In this version of the manuscript, some parts have been rearranged and edited for clarity; additional references have been added; and small corrections to the figures and text have been made.

